# Variance in within-pair reproductive success influences the opportunity for selection annually and over the lifetimes of males in a multi-brooded songbird

**DOI:** 10.1101/2020.03.03.974790

**Authors:** Ryan R. Germain, Michael T. Hallworth, Sara A. Kaiser, T. Scott Sillett, Michael S. Webster

**Affiliations:** Cornell Lab of Ornithology, Cornell University, Ithaca, NY, USA; Department of Neurobiology and Behavior, Cornell University, Ithaca, NY, USA; Department of Biology & GLOBE Institute, University of Copenhagen, Copenhagen, Denmark; Migratory Bird Center, Smithsonian Conservation Biology Institute, National Zoological Park, Washington, DC, USA; Northeast Climate Adaptation Science Center, University of Massachusetts Amherst, Amherst, MA, USA

## Abstract

In socially monogamous species, male reproductive success consists of ‘within-pair’ offspring produced with their socially-paired mate(s), and ‘extra-pair’ offspring produced with additional females throughout the population. Both reproductive pathways offer distinct opportunities for selection in wild populations, as each is composed of separate components of mate attraction, female fecundity, and paternity allocation. Identifying key sources of variance and covariance among these components is a crucial step towards understanding the reproductive strategies that males use to maximize fitness both annually and over their lifetimes. We use 16 years of complete reproductive data from a population of black-throated blue warblers (*Setophaga caerulescens*) to partition variance in male annual and lifetime reproductive success, and thereby identify if the opportunity for selection varies over the lifetimes of individual males and what reproductive strategies likely favor maximum lifetime fitness. The majority of variance in male reproduction was attributable to within-pair success, but the specific effects of individual components of variance differed between total annual and total lifetime reproductive success. Positive overall lifetime covariance between within-pair and extra-pair components indicates that males able to maximize within-pair success, particularly with double-brooding females, likely achieve higher overall lifetime fitness via both within-pair and extra-pair reproductive pathways.

## Introduction

Sexual selection represents the evolutionary processes that result from differences in mating or fertilization success within a population (Andersson 1994). The opportunity, or potential, for such selection to operate represents the ‘upper limit’ of selection in a population, and is proportional to the population-wide variance in mating and/or reproductive success; if variance is low (e.g. all individuals have similar numbers of mates and offspring) then selection will be weak, whereas if variance in reproductive output is high (e.g. few individuals monopolize mating opportunities and produce many offspring), selection potentially can be strong (Crow 1958, 1991; Emlen and Oring 1977; Wade and Arnold 1980; Jones 2009; Shuster 2009). However, an individual’s total reproductive success (*T*_*RS*_) is the result of several underlying sources of life-history variation, each of which may have different population-wide variances or may co-vary with different ecological and/or social factors, and hence represent distinct avenues through which selection may operate (Webster et al. 1995; Whittingham and Dunn 2004; Freeman-Gallant et al. 2005; Dolan et al. 2007; Lawler 2007; Lebigre et al. 2012). Determining where the greatest sources of variance and covariance exist among these life-history components is a key method to identifying the reproductive strategies that may enable individuals to maximize fitness annually and over their lifetimes (Kakauer et al. 2011).

Male *T*_*RS*_ is comprised of two pathways in socially monogamous yet genetically promiscuous species: ‘within-pair’ offspring produced by their social mate(s) (i.e. females with which they share reproductive duties such as territory defense and parental care), and additional ‘extra-pair’ offspring produced throughout the population by females socially paired to other males (Birkhead and Møller 1995; Webster et al. 1995; Whittingham et al. 2006). Both within-pair and extra-pair reproductive success (*W*_*RS*_, *E*_*RS*_) are themselves comprised of components of mate attraction (i.e. pre-copulatory) and paternity (i.e. post-copulatory) success, as well as by the fecundity of a male’s within-pair and extra-pair mates (Figure 1). Many studies have sought to identify sources of variance among mate attraction and paternity success in different taxa (e.g. Gibson and Guiness 1980; Lane et al. 2009; Vedder et al. 2011; Losdat et al. 2015; Evans and Garcia-Gonzalez 2016), but less emphasis has been placed on the importance of female fecundity in driving variation in male *T*_*RS*_. In particular, the ability of some females in a population to produce multiple broods or litters of offspring within a given breeding season has the potential to markedly increase overall variance in male *W*_*RS*_ by increasing the number of within-pair offspring they can potentially sire (e.g. Nagy and Holmes 2005b; Kaiser et al. 2017).

**Figure 1:**
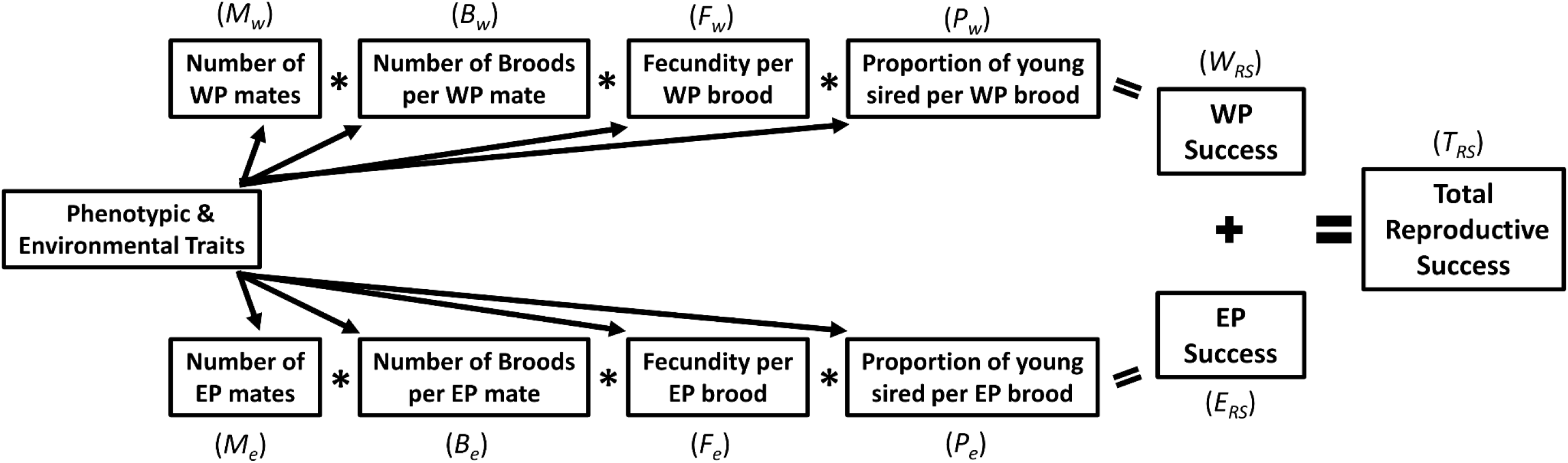
Conceptual diagram depicting the components of within-pair (WP) and extra-pair (EP) reproductive success, which combine to determine an individual male’s total reproductive success. Arrows indicate the effects that any potential environmental and/or phenotypic traits can have on WP and EP components (including the number of broods and fecundity per brood of his WP and EP female mates), and hence on total reproductive success. Italic terms (in parentheses) denote abbreviations used for each component of variance. Note that components of within-pair and extra-pair pathways are multiplicative, whereas WP and EP success themselves are additive components of total annual reproductive success (adapted from Webster et al. 1995).

Alternatively, if the extra-pair pathway is of primary importance for driving variance in total reproductive success in such populations, multi-brooding should not affect variance in *W*_*RS*_, but rather allow some males to capitalize on additional opportunities to gain extra-pair success with multi-brooding females throughout the population, leading to greater variance in *E*_*RS*_ than in *W*_*RS*_ (e.g. Webster et al. 2007; Lebigre et al. 2012; Losdat et al. 2015). Thus, quantifying the relative influence of component sources of within-pair and extra-pair female fecundity (i.e. number of broods or litters, number of offspring per brood or litter) on male *T*_*RS*_ has the potential to highlight the life-history traits which may influence independent opportunities for selection to operate in wild populations.

Much work has focused on the role that *E*_*RS*_ can play in driving the opportunity for selection in free-living, socially monogamous populations (e.g. Yezerinac et al. 1995; Dolan et al. 2007; Webster et al. 2007; Balenger et al. 2009; Vedder et al. 2011; Lebigre et al. 2013), while other authors suggest that many of the purported effects of *E*_*RS*_ on the opportunity for selection may be inflated by incomplete sampling of potential extra-pair sires (Whittingham and Dunn 2004; Freeman-Gallant et al. 2005; Albrecht et al. 2007), or may vary with annual variation in social-environmental factors such as population density (Møller and Birkhead 1993; Taff et al. 2013). However, the majority of field-based studies investigating the opportunity for selection are conducted over relatively short timescales and are thus limited to decomposing variance in annual reproductive success only. Given within-individual heterogeneity in annual reproduction, individual strategies to maximize annual or lifetime reproductive success may differ, depending on socio-environmental context or individual phenotype. If selection acted consistently over the lifetimes of individuals, we would observe little variation among years in the effects of individual components of *W*_*RS*_ and *E*_*RS*_ on total variance in annual reproductive success (*T*_*ARS*_). Similarly, under consistent selection, individual components of *W*_*RS*_ and *E*_*RS*_ would exhibit consistent effects on *T*_*ARS*_ and total variance in lifetime (*T*_*LRS*_) reproductive success, as well as realized mean annual and lifetime reproductive success in the population (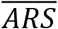 and 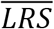, respectively). If, however, individual males trade off shorter-versus longer-term reproductive success, the effects of individual components of variance in *W*_*RS*_ and *E*_*RS*_ would exhibit opposing patterns between annual and lifetime reproductive success. Therefore, evaluating how components of reproduction that affect overall variance in *T*_*ARS*_ and *T*_*LRS*_ may differ, and quantifying how such components actually affect realized 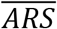 and 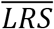, are crucial to our understanding of how the opportunity for selection operates in wild populations.

For males of many species, particularly those that engage in bi-parental care, reproductive gains through extra-pair mating may be offset by losses in within-pair paternity, resulting in a negative covariance (and hence a reproductive trade-off) between *W*_*RS*_ and *E*_*RS*_. For instance, seeking extra-pair copulations may come at the expense of losing within-pair paternity through reduced mate-guarding (Westneat and Stewart 2003; Kokko and Morrell 2005; Dias et al. 2009; Harts et al. 2016; Reitsma et al. 2018). In contrast, some studies find positive overall covariance between *W*_*RS*_ and *E*_*RS*_, indicating that males with a higher propensity for acquiring extra-pair mates and/or siring extra-pair young are also more likely to have higher within-pair success (Yezerinac et al. 1995; Albrecht et al. 2007; Ferree and Dickinson 2014; Reid et al. 2014). We currently have a poor understanding of where potential sources of such trade-offs exist in wild populations, in part due to the data requirements needed to accurately quantify covariance among individual components of *W*_*RS*_ and *E*_*RS*_. Co-variances have the potential to dramatically shape the evolutionary dynamics of reproductive systems (Lebigre et al. 2013; Reid et al. 2014). Evaluating sources of covariance among individual components of *W*_*RS*_ and *E*_*RS*_ is therefore a key facet of identifying potential sources of reproductive trade-offs and thus quantifying the overall opportunity for sexual selection to operate.

Here, we use 16 years of detailed breeding observations and genetic paternity data from a longitudinal study of black-throated blue warblers (*Setophaga caerulescens*) to identify the contributions of components of *W*_*RS*_ and *E*_*RS*_, as well as their covariances, to the opportunity for selection in this socially monogamous, multi-brooded songbird. We decompose *T*_*ARS*_ and *T*_*LRS*_ to their component sources of (co)variance to identify which sources represent the greatest overall opportunity for selection to operate in this system, both annually and over the lifetimes of individual males. Secondly, we test for the presence of reproductive trade-offs between components of *W*_*RS*_ and *E*_*RS*_. If strong sources of negative covariance exist among components of *W*_*RS*_ and *E*_*RS*_, it would suggest that selection favors males trading off one reproductive pathway for the other. If, however, there is non-negative covariance between *W*_*RS*_ and *E*_*RS*_, it would suggest that males which employ reproductive strategies that achieve higher success in one pathway experience higher overall fitness on average, and that selection favors males which are capable of capitalizing on multiple facets of reproductive success.

## Material and Methods

### Population monitoring and focus on selection in males

We monitored an individually marked population of black-throated blue warblers over 16 years (1999-2015) at the Hubbard Brook Experimental Forest (3,160ha) in New Hampshire, USA (43°56’N, 71°45’W). The breeding ecology of this long-term study population is described in detail elsewhere (e.g. Rodenhouse et al. 2003; Sillett et al. 2004; Holmes 2011; Kaiser et al. 2017). In brief, black-throated blue warblers are migratory, territorial songbirds found in relatively high densities at our study site. Arrival and the onset of breeding typically occurs in early May (Holmes et al. 2017), and males defend 1-4 ha territories until breeding ends in August (Sillett et al. 2004). Each season, adult males (socially-paired and unpaired) and females are captured in mist nets, marked with a unique combination of three colored leg bands and one aluminium U.S. Geological Survey (USGS) leg band, and blood sampled (~70 μl) from the brachial vein. All nesting attempts are monitored on three study plots (35-85 ha). Nestlings are banded with one USGS leg band and blood sampled (~30 μl) on day 6 after hatching, and then monitored daily until nest departure (i.e. fledge date, approx. 9 days after hatching; Holmes et al. 2017).

Our current study focused on the opportunity for selection in males, since sexual selection tends to operate more strongly among males than among females in many animal populations (Bateman 1948; Wade 1979; Arnold and Duvall 1994; Shuster 2009; but see Clutton-Brock 2007). Females are often the choosier sex because they are energetically limited in the number of offspring they can produce (i.e. fecundity) and show relatively little among-individual variation in reproductive success compared to males (e.g. Hoogland and Foltz 1982; Freeman-Gallant et al. 2005; Lebigre et al. 2012). In contrast, male reproductive success is often limited by the number of potential mates they can attract and by the number of offspring that they sire throughout the population, leading to greater variance in *T*_*RS*_ as some males attract numerous females and/or sire many offspring while others have low mating or siring success (e.g. Cerchio et al. 2005; Freeman-Gallant et al. 2005; Webster et al. 2007; Lebigre et al. 2012; Dubuc et al. 2014). Decomposing the potentially complex components of variance in female reproductive success is an important topic of consideration in its own right; however, such analyses are beyond the scope of our current study and, given lower variance compared to males, will likely be further informed by first identifying the components of female fecundity which may influence variation in male within-pair and extra-pair reproductive success (below).

### Genetic paternity assignment and calculation of annual/lifetime reproductive success

Detailed methods regarding genetic paternity assignment for this system are described by Kaiser et al. (2017). Briefly, we genotyped 4097 offspring and >95% of all adults in the study area (including candidate males adjacent to plot boundaries) at six highly polymorphic microsatellite loci using Genemapper v.4.1 (Applied Biosystems). We conducted parentage analyses for each year and study plot separately using CERVUS v3.0 (Kalinowski et al. 2007), which uses a maximum likelihood-based approach to infer parentage. The above methods resulted in a combined probability of paternal exclusion of 0.999 (Kaiser et al. 2017).

We define a male’s annual reproductive success as the total number of fledglings that he sired in a given year. Within-pair success represents the number of sired offspring fledged from nesting attempts by his social female(s) on his territory, while extra-pair success represents all other fledged offspring that he sired within the population that year. Lifetime reproductive success is defined as a male’s summed annual reproductive success over his lifetime as a breeder in the population. An individual male was classified as dead/no-longer breeding if they were no longer observed in the population (e.g. defending a territory or mating/caring for offspring) and if no further offspring were genetically assigned to the focal male in any subsequent year. Genetic paternity data were not available before 1999 and after 2016 at the time of analysis; therefore we excluded n = 11 males that were breeders before 1999 and n = 21 males that survived beyond 2016 from analyses of lifetime reproductive success to avoid underestimating this metric for males that may have produced offspring outside of the study period.

### Statistical analyses

All analyses were performed in R 3.5.1 (R Development Core Team 2018). Our initial dataset consisted of 1108 observations of male annual reproductive success (i.e. ‘male-years’) from 1999-2015. For all analyses, we excluded observations from males that produced either within-pair or extra-pair offspring with females on territories that were part of brood manipulation (n = 50) or food supplementation (n = 79) experiments. To ensure that our analyses included the most accurate information possible on extra-pair reproductive success, we also excluded males that had a higher likelihood of siring unsampled extra-pair offspring outside of the study area (n = 210 males with territories located within 210m [the equivalent of ~3 territory lengths] of the plot boundaries, Kaiser et al. 2017). Results remained qualitatively similar between the reduced and full dataset (e.g. variance decomposition results changed by <1% for total annual reproductive success).

We first decomposed population-wide variance in *T*_*ARS*_ and *T*_*LRS*_ over the entire study period to identify the components of variance in annual and lifetime reproductive success that present the greatest opportunity for selection to operate. Variance in total reproductive success (*T*_*RS*_, here representing either total annual [*T*_*ARS*_] or lifetime [*T*_*LRS*_] reproductive success) is the sum of variances of a male’s within-pair reproductive success (*W*_*RS*_) and extra-pair reproductive success (*E*_*RS*_) success, as well as 2× their associated covariance:

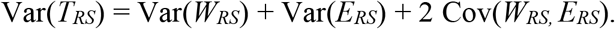

Following Webster et al. (1995), *T*_*RS*_ can be further partitioned into separate components of pairing success (i.e. number of mates, *M*), female fecundity, and paternity (*P*) allocation (and their associated covariances), which act multiplicatively to equal either *W*_*RS*_ or *E*_*RS*_ for an individual male (Figure 1). However, our variance decomposition differs from Webster et al. (1995) in that here we further partition female fecundity into two parts: the number of broods (*B*) produced per female, given that some females in our population will double-brood under favorable environmental conditions (Nagy and Holmes 2005a; Kaiser et al. 2015), and female fecundity per brood (i.e. the number of offspring produced per brood, *F*). For ease of interpretation, we present each component of variance or covariance as the population wide (co)variance of the component itself (e.g. *M*_*w*_ represents variance in within-pair pairing success, and Cov(*M*_*w*_,*M*_*e*_) represents the covariance between within-pair and extra-pair pairing success), but see Supporting Information S1 for full descriptions of how each variance component was calculated. Males that had zero success in one component of *W*_*RS*_ or *E*_*RS*_ were not included in analyses of downstream variance decomposition to avoid conflating variance estimates (following Webster et al. 1995; Freeman-Gallant et al. 2005). Note that this method of variance decomposition also estimates remainder terms, where the total remainder (*D*_*T*_) is equal to *D*_*w*_ + *D*_*e*_ + *D*_*we*_, which are each calculated by subtracting the summed variance of each individual component from the overall variance in *W*_*RS*_/*E*_*RS*_/their covariance; e.g. *D*_*w*_ = Var(*W*_*RS*_) – (Var[*M*_*w*_] + Var[*B*_*w*_] + Var[*F*_*w*_] + Var[*P*_*w*_] + Cov[*M*_*w*_, *B*_*w*_] + Cov[*M*_*w*_, *F*_*w*_] + Cov[*M*_*w*_, *P*_*w*_] + Cov[*B*_*w*_, *F*_*w*_] + Cov[*B*_*w*_, *P*_*w*_] + Cov[*F*_*w*_, *P*_*w*_]). These remainder terms capture multivariate skewness expressed in the higher order moments of the distributions of each component of reproductive success, such that total variance in *T*_*ARS*_ and *T*_*LRS*_ is not a simple sum of the component variances and covariances (Bohrnstedt and Goldberger 1969; Webster et al. 1995; Dolan et al. 2007; Lawler 2007).

We identified components of (co)variance in both *T*_*ARS*_ and *T*_*LRS*_ that constituted a substantial percentage (arbitrary cut-off of ≥10%) of the total variance across the full study period. For *T*_*ARS*_, we further conducted time-series analyses to evaluate whether these key components in annual reproductive success consistently provided the greatest opportunity for sexual selection in each year of study (i.e. under varying socio-environmental conditions such as population density or population age-structure). To do so, we replicated our variance decomposition (above) for each individual year in the study period, extracted the percent of total variance in *T*_*ARS*_ attributed to each key component, as well as overall (co)variance in *W*_*RS*_ and *E*_*RS*_, and both visually inspected time series plots for presence of lags and conducted Kwiatkowki-Phillips-Schmidt-Shin (KPSS) tests to determine if each time series was trend stationary (i.e. lack of overall significant positive/negative trend over the study period).

Lastly, we determined the relative effect of each key component of *W*_*RS*_ and *E*_*RS*_ on mean annual and lifetime reproductive success (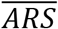 and 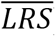) in the population using generalized mixed effects models and a model averaging approach. For 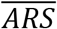, we constructed a global mixed model (Poisson distribution) which included male identity and year as random effects, and *M*_*w*_, *B*_*w*_, *P*_*w*_, *M*_*e*_, *B*_*e*_ as fixed effects (see Table 1 for justification) as well as the interaction between *B*_*w*_ and *B*_*e*_, to represent the covariance between these two components that explain a substantial portion of variance in *T*_*ARS*_ (Table 1). All fixed effects were standardized to mean = 0, SD = 1 to reduce any influence of measurement scale on model results and to allow direct comparison of model coefficients (White and Burnham 1999). We ran all possible combinations of these fixed effects (n = 40 models total) and selected a subset with a difference in Akaike Information Criterion (ΔAIC) ≤ 7 from the best-fitting model. We chose a cutoff of ΔAIC ≤ 7 to ensure that estimates of the relative influences of each predictor on 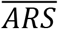 were as conservative and inclusive as possible (Burnham and Anderson 2002). We then averaged parameter estimates for each predictor included in this subset of models to create one representative (full-average) estimate of the relative effects of each component on 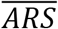 (Burnham and Anderson 2002; Germain and Arcese 2014; Germain et al. 2015, 2018). Statistical significance for each fixed effect was assessed by whether 95% confidence intervals (CIs) overlapped zero.

**Table 1:**
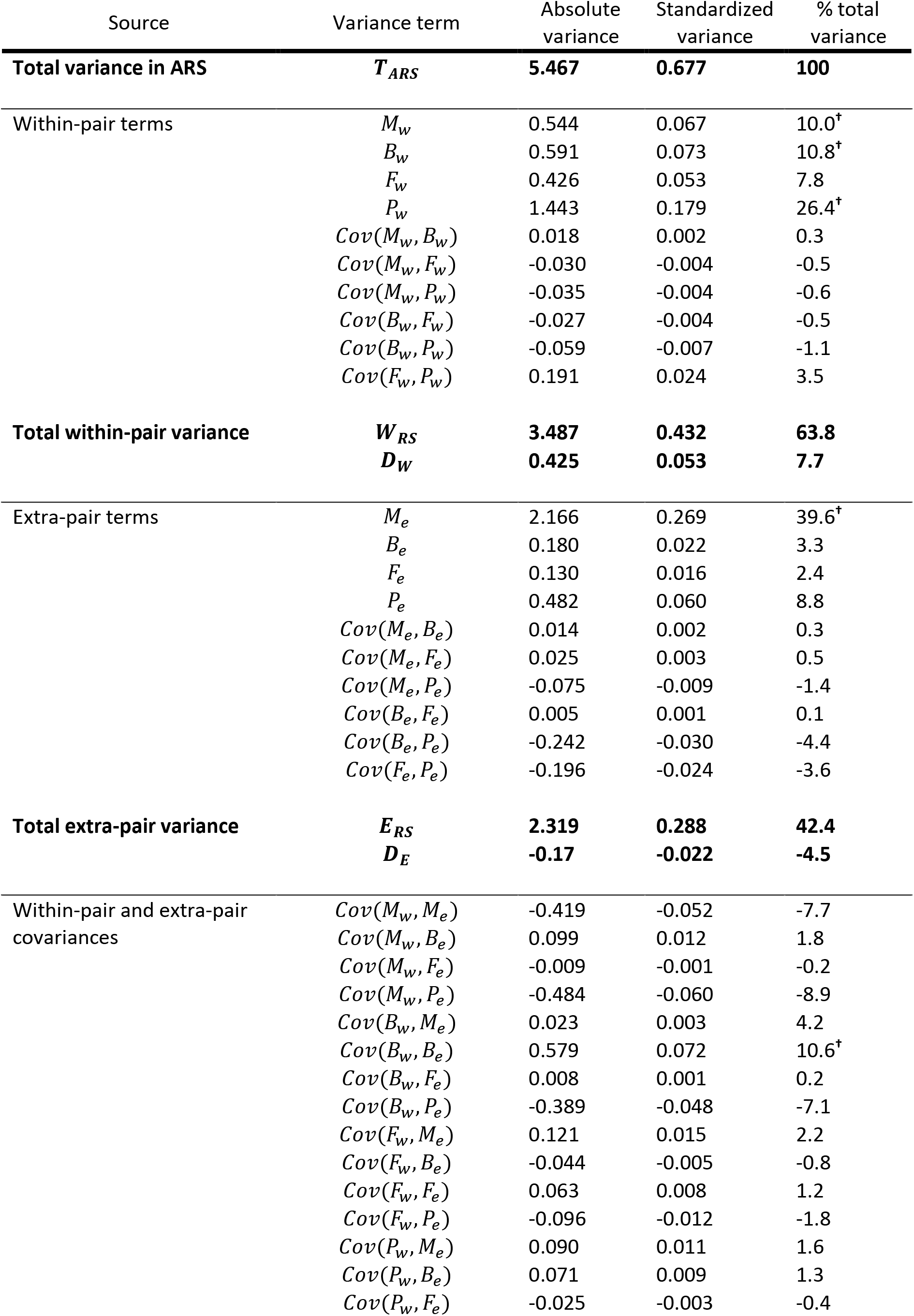
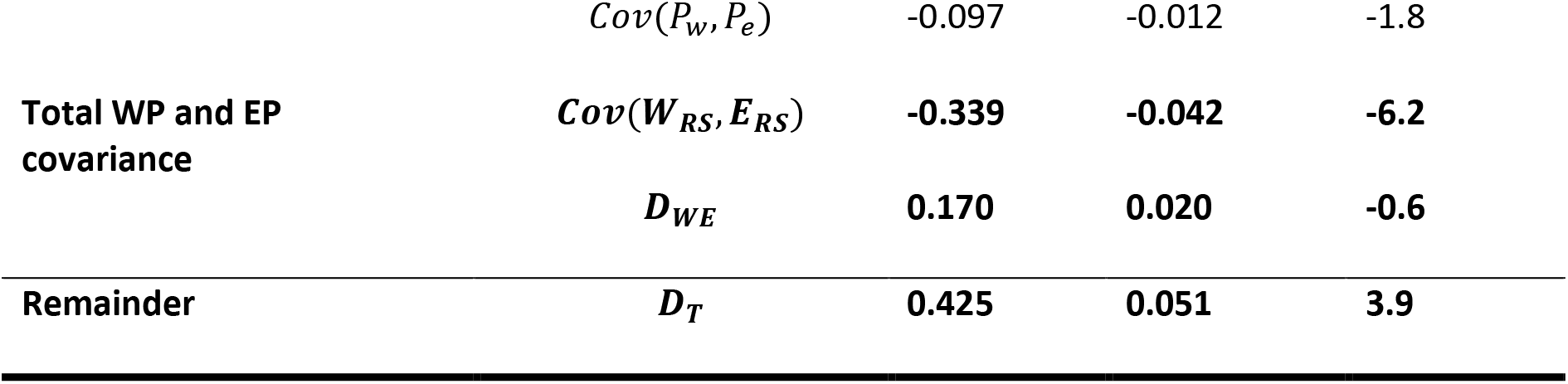
Decomposition of variance in total annual reproductive success (*T*_*ARS*_) of black-throated blue warblers across 16 years of study. For each row, variance terms (see Figure 1 for definitions) represent the focal component of (co)variance in *T*_*ARS*_, but note that actual calculation of each term involves population-wide means for additional components (see Supporting Information S1 and Webster et al. 1995 for further details). *D* denotes remainder terms for within-pair, extra-pair, and covariance pathways, where *D*_*T*_ = *D*_*w*_ + *D*_*e*_ + *D*_*we*_. Values depict the absolute variance in each component, as well as standardized variance (i.e. divided by population mean 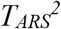 = 8.713), and the percentage of total variance in *T*_*ARS*_ attributable to each term (i.e. divided by Var[*T*_*ARS*_]). Crosses (^**†**^) denote variance terms which fall above our cutoff of contributing >10% of total variance in *T*_*ARS*_.

We repeated this approach for 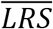, constructing a global mixed model (Poisson distribution) which included male identity and male longevity (years breeding in the population) as random effects, and *M*_*w*_, *B*_*w*_, *F*_*w*_, *P*_*w*_, *M*_*e*_, *B*_*e*_, *F*_*e*_, and *P*_*e*_ as fixed effects, as well as the interactions between *M*_*w*_ × *F*_*w*_, *M*_*e*_ × *F*_*e*_, *M*_*w*_ × *M*_*e*_, *M*_*w*_ × *F*_*e*_, *M*_*w*_ × *P*_*e*_, *B*_*w*_ ×*B*_*e*_, *B*_*w*_ × *P*_*e*_, *F*_*w*_ × *M*_*e*_, and *F*_*e*_ × *F*_*w*_ to represent covariances among these key components of variance in *T*_*LRS*_ (see Table 2 for justification). We again ran all possible combinations of these fixed effects (n = 4458 models), selected those within ΔAIC ≤ 7 of the best-fitting model, and averaged parameter estimates for each predictor included in this subset (as above).

**Table 2:** Decomposition of variance in total lifetime reproductive success (*T*_*LRS*_) of black-throated blue warblers across 16 years of study. For each row, variance terms (see Figure 1 for definitions) represent the focal component of (co)variance in *T*_*LRS*_, but note that actual calculation of each term involves population-wide means for additional components (see Supporting Information S1 and Webster et al. 1995 for further details). *D* denotes remainder terms for within-pair, extra-pair, and covariance pathways, where *D*_*T*_ = *D*_*w*_ + *D*_*e*_ + *D*_*we*_. Values depict the absolute variance in each component, as well as standardized variance (i.e. divided by population mean 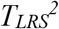 = 20.57), and the percentage of total variance in *T*_*LRS*_ attributable to each term (i.e. divided by Var[*T*_*LRS*_]). Crosses (^**†**^) denote variance terms which fall above our cutoff of >10% total variance in *T*_*LRS*_.

| Source | Variance term | Absolute variance | Standardized variance | % total variance |
| --- | --- | --- | --- | --- |
| **Total Variance** | **$T_{LRS}$** | **21.77** | **1.058** | **100** |
| Within-pair terms | $M_w$ | 5.766 | 0.280 | 26.5<sup>†</sup> |
| | $B_w$ | 5.331 | 0.259 | 24.5<sup>†</sup> |
| | $F_w$ | 7.672 | 0.373 | 35.2<sup>†</sup> |
| | $P_w$ | 6.140 | 0.299 | 28.2<sup>†</sup> |
| | $Cov(M_w, B_w)$ | 1.718 | 0.084 | 7.8 |
| | $Cov(M_w, F_w)$ | 12.383 | 0.602 | 56.9<sup>†</sup> |
| | $Cov(M_w, P_w)$ | 0.539 | 0.026 | 2.5 |
| | $Cov(B_w, F_w)$ | -0.213 | -0.010 | -1.0 |
| | $Cov(B_w, P_w)$ | -0.165 | -0.008 | -0.8 |
| | $Cov(F_w, P_w)$ | 0.908 | 0.044 | 4.2 |
| **Total within-pair variance** | **$W_{RS}$** | **10.171** | **0.494** | **46.7** |
|  | **$D_W$** | **-29.908** | **-1.455** | **-137.3** |
| Extra-pair terms | $M_e$ | 19.627 | 0.954 | 90.1<sup>†</sup> |
| | $B_e$ | 1.216 | 0.059 | 5.6 |
| | $F_e$ | 7.345 | 0.357 | 33.7<sup>†</sup> |
| | $P_e$ | 3.336 | 0.162 | 15.3<sup>†</sup> |
| | $Cov(M_e, B_e)$ | 0.318 | 0.016 | 1.5 |
| | $Cov(M_e, F_e)$ | 21.767 | 1.047 | 99.9<sup>†</sup> |
| | $Cov(M_e, P_e)$ | -1.491 | -0.072 | -6.8 |
| | $Cov(B_e, F_e)$ | 0.259 | 0.013 | 1.2 |
| | $Cov(B_e, P_e)$ | -1.752 | -0.085 | -8.1 |
| | $Cov(F_e, P_e)$ | -1.773 | -0.086 | -8.1 |
| **Total extra-pair variance** | **$E_{RS}$** | **5.806** | **0.282** | **26.7** |
|  | **$D_E$** | **-43.046** | **-2.083** | **-197.6** |
| Within-pair and extra-pair covariances | $Cov(M_w, M_e)$ | 12.860 | 0.625 | 59.1<sup>†</sup> |
| | $Cov(M_w, B_e)$ | 0.394 | 0.019 | 1.8 |
| | $Cov(M_w, F_e)$ | 8.264 | 0.402 | 37.9<sup>†</sup> |
| | $Cov(M_w, P_e)$ | -3.141 | -0.145 | -14.4<sup>†</sup> |
| | $Cov(B_w, M_e)$ | 0.478 | 0.023 | 2.2 |
| | $Cov(B_w, B_e)$ | 3.648 | 0.177 | 16.8<sup>†</sup> |
| | $Cov(B_w, F_e)$ | 0.389 | 0.019 | 1.8 |
| | $Cov(B_w, P_e)$ | -2.627 | -0.128 | -12.1<sup>†</sup> |
| | $Cov(F_w, M_e)$ | 15.835 | 0.770 | 72.7<sup>†</sup> |
| | $Cov(F_w, B_e)$ | -0.432 | -0.021 | -2.0 |
| | $Cov(F_w, F_e)$ | 10.395 | 0.505 | 47.7<sup>†</sup> |
| | $Cov(F_w, P_e)$ | -0.319 | -0.015 | -1.5 |
| | $Cov(P_w, M_e)$ | 1.101 | 0.053 | 5.1 |
| | $Cov(P_w, B_e)$ | -0.166 | -0.008 | -0.8 |
| | $Cov(P_w, F_e)$ | 0.379 | 0.018 | 1.7 |
| | $Cov(P_w, P_e)$ | -0.484 | -0.024 | -2.2 |
| Total WP and EP covariance | $Cov(W_{RS}, E_{RS})$ | 5.792 | 0.278 | 26.6 |
| | $D_{WE}$ | -40.782 | -1.991 | -187.2 |
| Remainder | $D_T$ | -113.0736 | -5.529 | -522.1 |

## Results

Male black-throated blue warblers exhibited considerable variation in both within-pair (mean = 1.7 young sired ±1.9SD, range = 0–9) and extra-pair (mean = 1.1 young sired ±1.5SD, range = 0–9) annual reproductive success, leading to substantial variation in the total number of offspring produced annually (mean = 2.8 ±2.3SD, range = 0–14; Figure 2a). Similarly, males varied in the number of within-pair (mean = 2.76 ±3.2SD, range = 0–23) and extra-pair (mean = 1.77±2.4SD, range = 0–14) offspring produced over their lifetimes. Hence, the Hubbard Brook population had relatively large variance in lifetime reproductive success (mean = 4.54 ± 4.7SD, range = 0–28), where most males produced few offspring and a small proportion produced many (Figure 2b).

**Figure 2:**
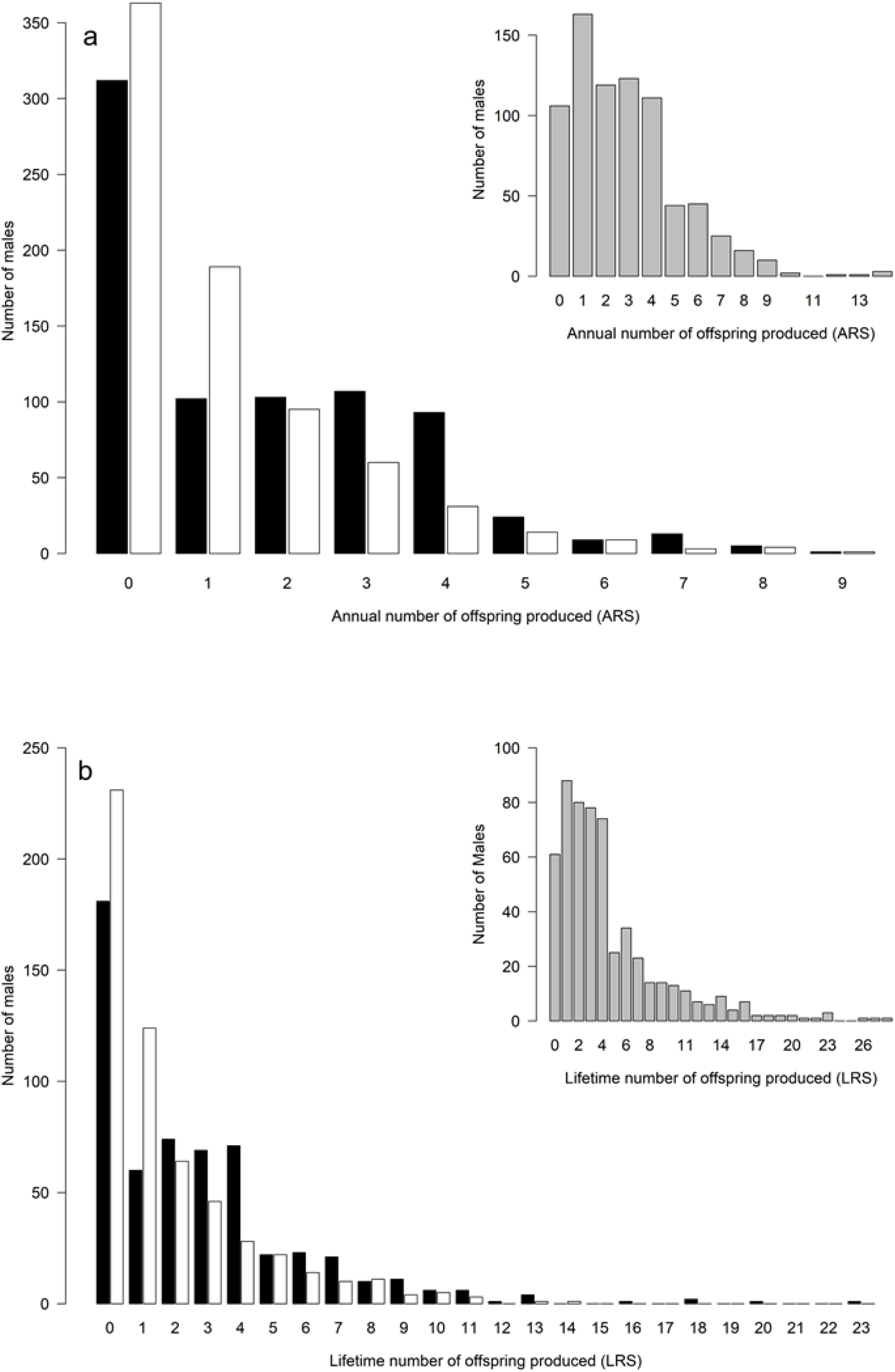
Total number of within-pair (black bars) and extra-pair (white bars) offspring produced by male black-throated blue warblers a) annually (n = 769 male-years) and b) over their complete breeding lifetime (n = 572 males). In each panel, insert (grey bars) depicts total number of offspring (within-pair and extra-pair) produced by these same males.

Within-pair and extra-pair components likewise varied throughout the population. Males had a higher mean number of within-pair mates (mean = 0.94 ± 0.41SD, range = 0–2) but exhibited a broader range in their number of extra-pair mates (mean = 0.69 ± 0.79SD, range = 0–5) within a given year. While the number of broods produced by within-pair (mean = 1.12 ± 0.51SD, range = 0–2) and extra-pair females (mean = 1.35 ± 0.45SD, range = 0–2) were similar, fecundity per brood was slightly lower for within-pair females (mean = 2.8 ± 1.08SD, range = 0–4) than extra-pair females (mean = 3.02 ± 0.85SD, range = 1–4.75), but note that sampling methodology requires a male to successfully sire one offspring to be included in analysis of extra-pair success. This is also reflected in the slight difference in range between the proportion of offspring sired per within-pair (mean = 0.58 ± 0.41SD, range = 0–1) and extra-pair (mean = 0.46 ± 0.25SD, range = 0.11–1) brood.

### The opportunity for selection in Annual Reproductive Success

Decomposition of the total opportunity for selection in annual reproductive success revealed that the majority of variance (~64%) in *T*_*ARS*_ was attributable to variance calculated from the within-pair pathway, of which within-pair paternity success (*P*_*w*_) constituted the greatest proportion (26.4%, Table 1). Both the number of within-pair mates (*M*_*w*_) and number of broods produced by within-pair mates (*B*_*w*_) also each accounted for roughly 10% of *T*_*ARS*_, suggesting multiple distinct components of selection within the *W*_*RS*_ pathway of annual reproductive success (Table 1). In contrast, the ~42% of variance in *T*_*ARS*_ attributable to the extra-pair pathway (*E*_*RS*_) was almost entirely accounted for by the number of extra-pair mates the male acquired (*M*_*e*_, 39.6%; Table 1). Overall covariance between *W*_*RS*_ and *E*_*RS*_ was slightly negative (−6.2% of total variance in *T*_*ARS*_), with only the positive covariance between the number of broods by within-pair and extra-pair females (Cov[*B*_*w*_, *B*_*e*_]) accounting for a substantial proportion of variance in *T*_*ARS*_ (10.6%, Table 1).

Variance in *T*_*ARS*_ varied dramatically across the 16-year study, indicating that the overall opportunity for selection differed among years (Figure 3a). Despite this, variance in *T*_*ARS*_ was trend stationary (KPSS trend = 0.13, p = 0.08), indicating no significant trend in the change in variance over the study period. The percent of *T*_*ARS*_ attributable to variance in *W*_*RS*_ likewise varied considerably, but also remained trend stationary (KPSS trend = 0.13, p = 0.07) and accounted for more total variance than *E*_*RS*_ in each year of study (Figure 3b). Among components of variance in within-pair reproductive success (Figure 3b), both number of within-pair mates (*M*_*w*_) and number of within-pair broods (*B*_*w*_) alternated above and below 10% of variance in *T*_*ARS*_, and while *M*_*w*_ exhibited no obvious trend (KPSS trend = 0.11, p = 0.1), variance in *B*_*w*_ exhibited a non-significant negative trend over the study period (KPSS trend = 0.15, p = 0.05, Figure 3b). Similarly, the proportion of within-pair offspring sired (*P*_*w*_), which consistently accounted for the largest percentage of *T*_*ARS*_ among within-pair components, also exhibited a non-significant negative trend (KPSS trend = 0.14, p = 0.05), due to high within-pair variance during the early portion of the study. Both *E*_*RS*_ and its main component, the number of extra-pair mates acquired, consistently accounted for ~40% of *T*_*ARS*_ and both were trend stationary (*E*_*RS*_ – KPSS trend = 0.05, p = 0.1; *M*_*e*_ – KPSS trend = 0.10, p = 0.1). In contrast, the overall covariance between *W*_*RS*_ and *E*_*RS*_ varied from −73% to 28% of total variance in *T*_*ARS*_ over the study period, where negative values indicate a trade-off between within-pair and extra-pair reproductive success, and which may counter-act high proportions of variance ascribed to *W*_*RS*_ or *E*_*RS*_ in particular years (e.g., 2000, 2001; Figure 3d). Despite this wide range of variance in *T*_*RS*_ attributed to *Cov*(*W*_*RS*_, *E*_*RS*_), the overall covariance between within-pair and extra-pair reproductive success remained trend stationary (KPSS trend = 0.07, p = 0.1), as did its main constituent component, the covariance between within-pair and extra-pair number of broods (KPSS test = 0.08, p = 0.1).

**Figure 3:**
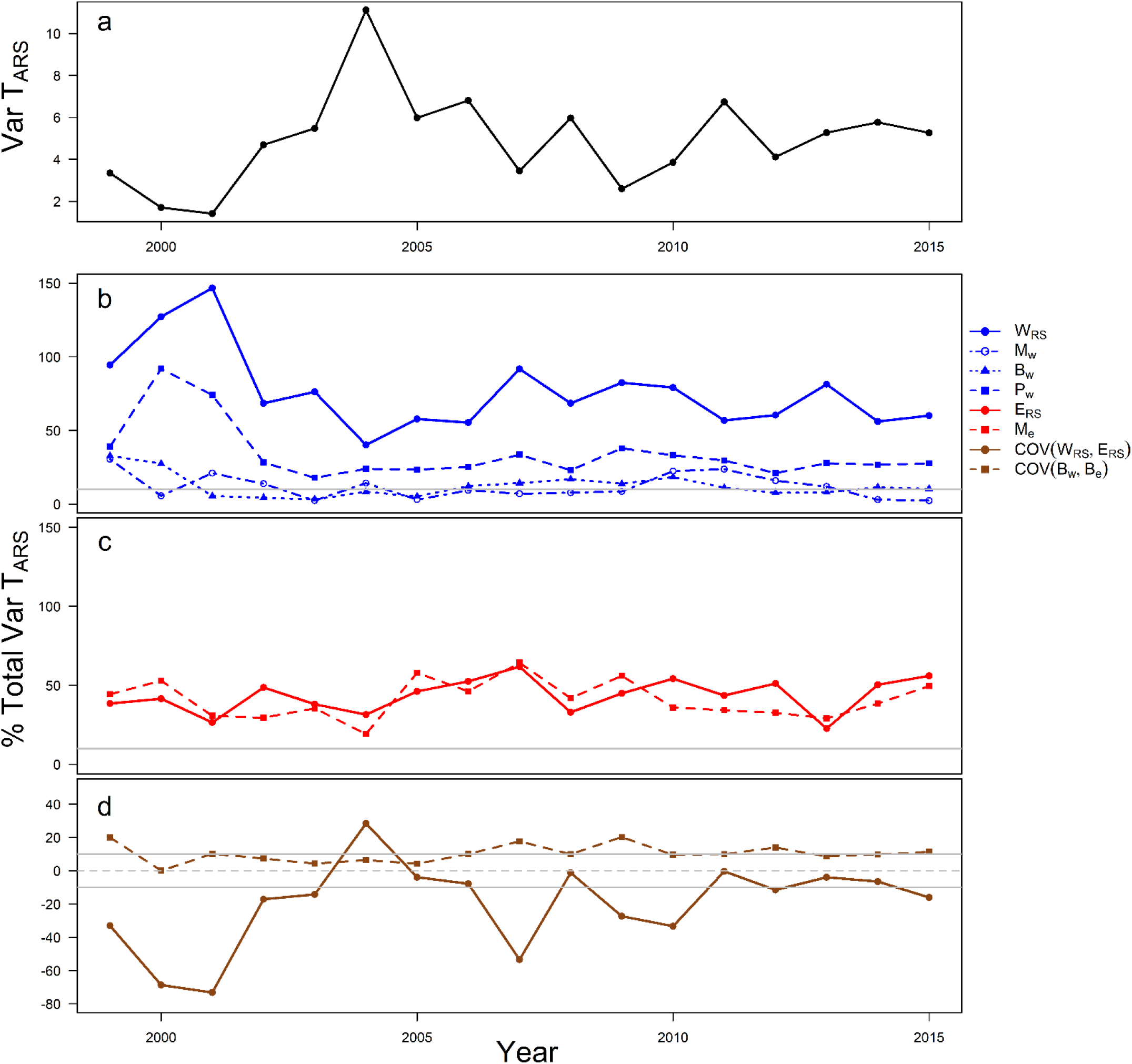
Time series plots depicting a) yearly variance in total annual reproductive success (*T*_*ARS*_) across a population of black-throated blue warblers, and yearly decompositions of the percent of total variance in *T*_*ARS*_ attributable to b) within-pair reproductive success (*W*_*RS*_), c) extra-pair reproductive success (*E*_*RS*_), and d) the covariance between within-pair and extra-pair reproductive success [*Cov(W*_*RS*_, *E*_*RS*_)]. For panels b, c, and d, colored dashed, dotted, and dot-dashed lines represent the yearly percentage of total variance in *T*_*ARS*_ attributable to constituent components of *W*_*RS*_, *E*_*RS*_, and *Cov(W*_*RS*_, *E*_*RS*_) found to represent ‘substantial variance’ (≥ 10%) across the study period (see Table 1), depicted as solid grey lines. For panel d, dashed gray line represents the transition from positive to negative covariance. Note different y-axis scales for panels a) and d).

Our global model (i.e. including all investigated predictor variables) of the effects of each key component on 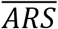 was a good fit to the data (R^2^ = 0.43). Our model subset included six models with ΔAIC ≤ 7 from the best-fitting model. All five fixed effects and the interaction between *B*_*w*_ and *B*_*e*_ were included in our final averaged model. Of these, *M*_*w*_, *B*_*w*_, *P*_*w*_, and *M*_*e*_ had significant, positive effects on 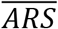 (Figure 4), indicating that males achieved the highest annual reproductive success by pairing with multiple double-brooded females, securing within-pair paternity, and mating with multiple extra-pair females (See Supporting Information S2 for full parameter estimates from final averaged model and all models in top subset).

**Figure 4:**
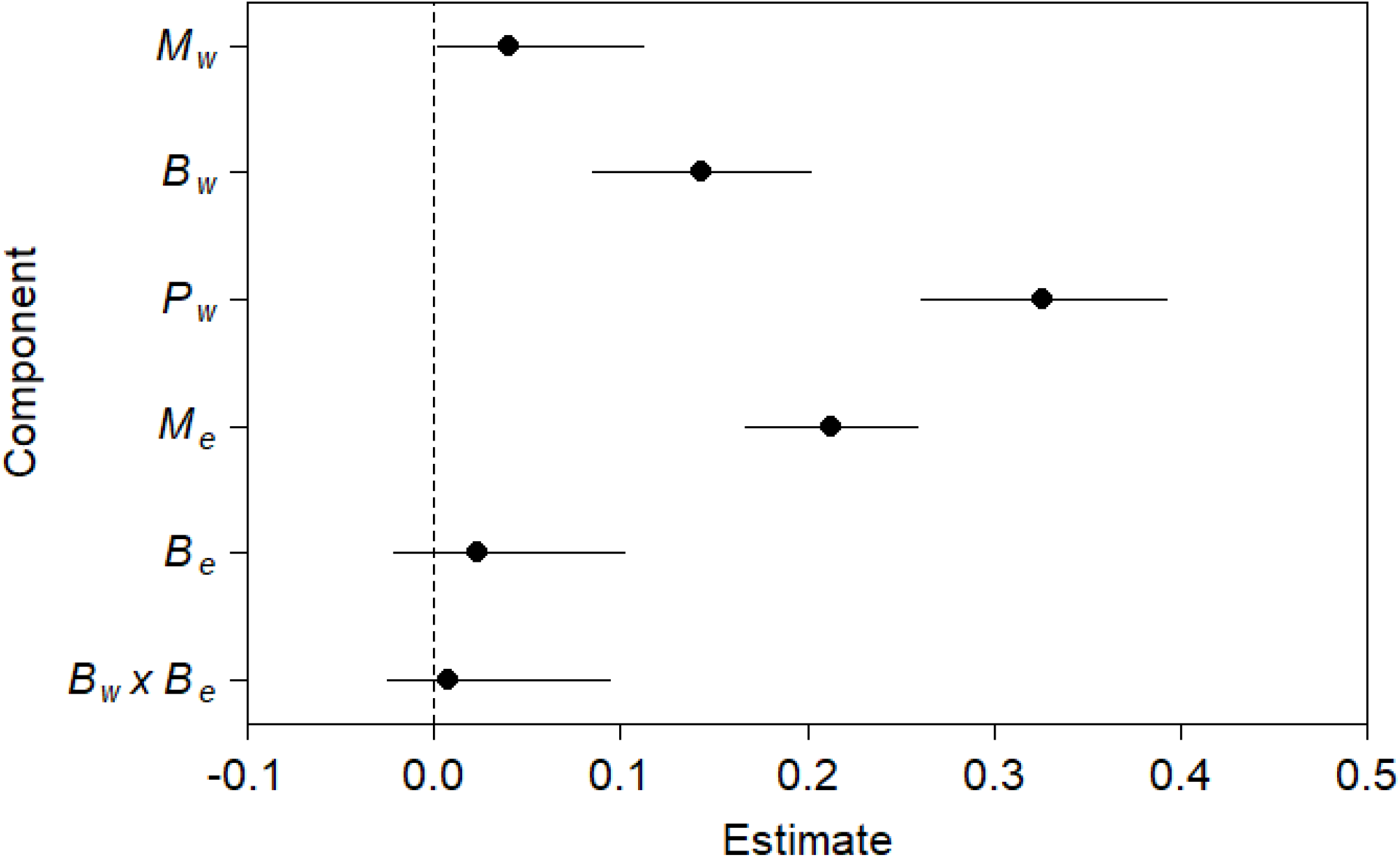
Effects plots from averaged generalized mixed-effects models testing the relative influence of each key component of (co)variance on mean annual reproductive success 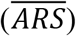. Points represent parameter estimates from final averaged model while whiskers depict 95% CIs. Dotted line represents 0, and parameter are considered significant if CIs do not overlap zero. *M*, *B*, and *P* refer to variance in number of mates, number of broods per mate, and proportion of young sired per brood for within-pair (*w*) or extra-pair (*e*) mates, respectfully. Full results from averaged model and all models in top subset are provided in Supporting Information S2.

### The opportunity for selection in Lifetime Reproductive Success

Decomposition of the opportunity for selection in lifetime reproductive success revealed that, similar to *T*_*ARS*_, substantially more variance in *T*_*LRS*_ was attributable to variance in within-pair reproductive success than to variance in extra-pair success (46.7% versus 26.7%, respectively, Table 2). Indeed, every individual component of *W*_*RS*_ contributed a substantial (>10%) proportion of total variance in *T*_*LRS*_, and the covariance between *M*_*w*_ and *F*_*w*_ accounted for over 50%. Similarly, although *M*_*e*_ again contributed the most of any single component to variance in *E*_*RS*_, *F*_*e*_ and *P*_*e*_ also accounted for ≥10% of total variance. The covariance between *M*_*e*_ and *F*_*e*_ contributed to almost 100% of variance in *T*_*LRS*_, indicating that the interaction between these two components is a strong driver of variance in male lifetime reproductive success (Table 2). Several components of within-pair/extra-pair covariances contributed to more than 10% of the total positive or negative variance in *T*_*LRS*_, suggesting the potential for both synergistic and trade-off effects between individual components of each pathway. However, overall covariance between *W*_*RS*_ and *E*_*RS*_ was positive and accounted for as much total variance in *T*_*LRS*_ as *E*_*RS*_ itself (~27%), indicating that there is no overall trade-off between the within-pair and extra-pair pathways in terms of male lifetime reproductive success in this system. Despite the strong potential for these listed sources of (co)variance to affect overall population-wide variance in male *T*_*LRS*_, large, negative sources of (co)variance (*D*_*w*_, *D*_*e*_, *D*_*we*_, *D*_*T*_, Table 2) were unaccounted for in our decomposition. Although the biological significance of such remainder terms is difficult to interpret without knowledge of how higher-order moments of the distribution of each within-pair and extra-pair component affect lifetime reproductive success (e.g. multivariate skewness: Bohrnstedt and Goldberger 1969; Lawler 2007), their relatively large, negative values suggest that some amount of realized variation in *T*_*LRS*_ is not captured by decomposing variance to its within-pair and extra-pair components and their associated covariances.

Our global model of the effects of each key component on mean lifetime reproductive success was an extremely good fit to the data (R^2^ = 0.86). A total of 147 models were included in the subset of models within ΔAIC ≤ 7 from the best-fitting model, which included all fixed effects and all interaction terms. We detected significant, positive effects of *B*_*w*_, *F*_*w*_, *P*_*w*_, *B*_*e*_, *F*_*e*_, *P*_*e*_ on 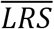, and significant negative effects from the interactions between *M*_*w*_ x *F*_*w*_, and *M*_*e*_ x *F*_*e*_, indicating that males in this population achieve the highest lifetime reproductive success by pairing with fewer double-brooded females that produced more offspring per brood and allocated more within-pair paternity to the focal male, and by mating with fewer double-brooding extra-pair females that likewise produced more offspring per brood and allocated more paternity towards the focal male. All other terms had CIs that overlapped zero (Figure 5), and were not considered to have statistically significant effects on 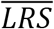 (see Supporting Information S3 for full parameter estimates from final averaged model and all models in top subset).

**Figure 5:**
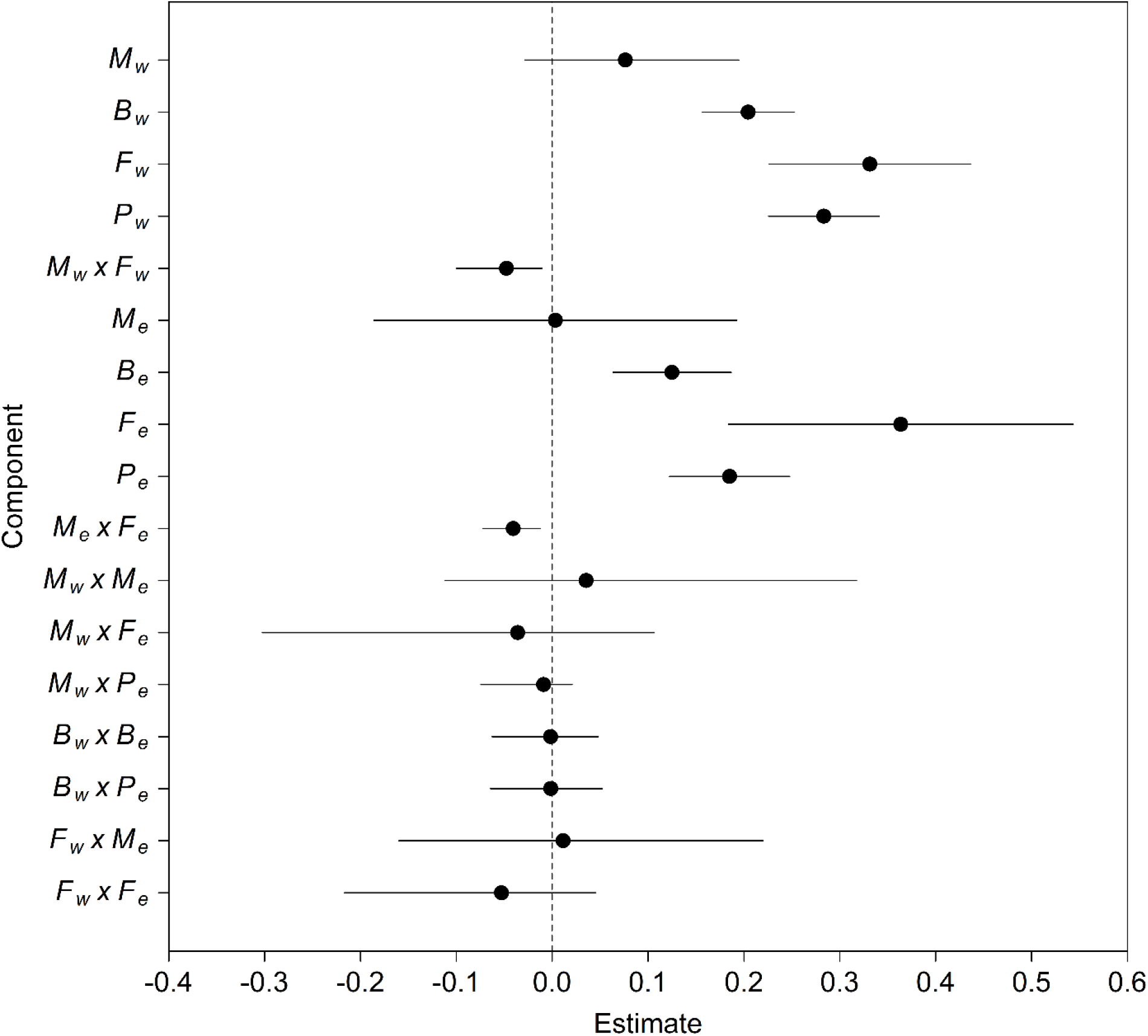
Effects plots from averaged generalized mixed-effects models testing the relative influence of each key component of (co)variance on mean lifetime reproductive success (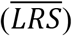). Points represent parameter estimates from final averaged model while whiskers depict 95% CIs. Dotted line represents 0, and parameters are considered significant if CIs do not overlap zero. *M*, *B*, *F*, and *P* refer to variance in number of mates, number of broods per mate, fecundity per brood, and proportion of young sired per brood for within-pair (*w*) or extra-pair (*e*) mates, respectfully. Full results from averaged model and all models in top subset are provided in Supporting Information S3.

## Discussion

Quantifying the opportunity for selection among different components of individual reproductive success is a key method for determining the evolutionary dynamics of mating systems within wild animal populations (Webster et al. 1995; Reid et al. 2014; Losdat et al. 2015). By identifying where the greatest opportunities for selection exist among various components of reproduction, and determining how the effects of these components may change annually or over the lifetimes of individual males, we can elucidate the reproductive strategies that males employ to maximize reproductive success as well as determine how such strategies may change over time. In a free living, multi-brooded songbird population, we found that among-individual variation in within-pair components of reproductive success consistently accounted for more of the total overall variance in both annual and lifetime reproductive success than did extra-pair components of variance (Table 1, Table 2, Figure 3). Several of these within-pair components were likewise associated with higher mean annual (Figure 4) and lifetime (Figure 5) reproductive success, and generally exhibited non-negative covariance with extra-pair components, suggesting that allocating resources towards attracting and defending access to socially-paired mates is likely a key tactic for maximizing annual and lifetime fitness in species which exhibit environmentally-mediated multiple brooding.

The majority of variance in total annual reproductive success (*T*_*ARS*_) was accounted for by four individual components of variance and one component of covariance which each explained ≥10% (Table 1). Although the percentage of *T*_*ARS*_ accounted for by *W*_*RS*_ did vary somewhat from year-to-year in our study (Figure 3b), the influences of each individual component of *W*_*RS*_ were relatively stable over the longer-term, indicating that individual behavioral strategies aimed at maximizing annual reproductive success are likely under consistent selection in this population. This finding contrasts with previous work indicating that the influence of particular variance components on the opportunity for selection can co-vary with socio-environmental factors like annual breeding density over shorter time periods (e.g. Møller and Birkhead 1993; Taff et al. 2013; Evans and Garcia-Gonzalez 2016). The overall covariance between *W*_*RS*_ and *E*_*RS*_ was slightly negative for *T*_*ARS*_ (Table 1) and tended to be negative in most years (Figure 3d); however, the positive covariance between the number of broods produced by a male’s socially-paired mate(s) and his extra-pair mate(s) (Cov[*B*_*w*_, *B*_*e*_]) was relatively consistent in accounting for ≥10% of total variance in *T*_*ARS*_, and was never negative over the 16 year study (Figure 3d). This indicates that while the opportunity for selection to operate on synergistic effects of multiple brooding among within-pair and extra-pair mates can vary somewhat annually, males never experienced a trade-off between the number of broods their socially-paired and extra-pair mates produced. While future studies will need to pinpoint the precise spatial and/or environmental variables which drive the positive covariance between within-pair and extra-pair double brooding, our results indicate that this covariance makes a relatively small but important contribution towards driving the opportunity for selection in this system.

In practice, all four key individual components of variance in *T*_*ARS*_ (i.e. *M*_*w*_, *B*_*w*_, *P*_*w*_, and *M*_*e*_) had significant, positive effects on mean annual reproductive success (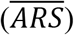) in our study population (Figure 4). Previous work from this population likewise indicates that greater investment in aspects of within-pair reproduction results in a higher reproductive net-gain for individual males, and that ‘high quality’ males on food-rich territories tend to invest in behaviors that enhance *W*_*RS*_ (e.g. increased mate-guarding and territorial defense) over those that enhance *E*_*RS*_ (e.g. extra-territorial forays; Kaiser et al. 2015, 2017). Food limitation appears to be the proximate mechanism behind such territorially-driven variation in mating strategies at Hubbard Brook, as females nesting in food-rich territories are more likely to double brood than those occupying food-limited habitat, with little loss in terms of individual energetic balance or lifespan (Holmes et al. 1996; Nagy and Holmes 2005a; Kaiser et al. 2015, 2017; Lany et al. 2016;). Taken together, our results indicate that securing a high-quality breeding territory is the key strategy for males in this population to maximize annual reproductive success via the within-pair pathway, and that doing so does not necessarily limit their ability to also achieve some *E*_*RS*_ through mating with multiple extra-pair females. Indeed, doing so may occasionally result in a higher reproductive net gain via the positive covariance between within-pair and extra-pair female brooding (Table 1, Figure 3). With warming temperatures, earlier springs and longer breeding seasons increase the frequency of double-brooding in our study species and other opportunistically double-brooding passerine birds (Townsend et al. 2013; Both et al. 2019), which may explain the near-significant negative trend in *B*_*w*_ over our study (i.e. more double-brooding females will decrease overall variance in *B*_*w*_; Figure 3). Increased growing season length with earlier leaf-out date and later leaf senescence, documented at Hubbard Brook (Richardson et al. 2006) and elsewhere (Ibáñez Inés et al. 2010; Fridley 2012) could therefore translate to higher realized annual reproductive success for males able to capitalize on both maximizing *W*_*RS*_ among their double-brooding socially-paired mate(s) and gaining additional *E*_*RS*_ with extra-pair females throughout the population.

Within-pair components of variance also accounted for the majority of variance in total lifetime reproductive success (*T*_*LRS*_), as we found for *T*_*ARS*_, but the relative influences of within-pair and extra-pair sources of (co)variance on the opportunity for selection differed somewhat between annual and lifetime reproduction (Table 2). Further, more individual components of *T*_*LRS*_ accounted for ≥10% of variance than those of *T*_*ARS*_. Several of these individual components accounted for over half (to almost all) of the total variance in *T*_*LRS*_ before taking negative covariance terms and remainder (*D*) terms into account (Table 2). Indeed, a striking feature of our variance decomposition of *T*_*LRS*_ is that several covariance terms, most notably between individual within-pair and extra-pair components, accounted for substantial proportions of variance, with the overall covariance between *W*_*RS*_ and *E*_*RS*_ accounting for as much of the total variance as overall extra-pair variance (Table 2). This highlights that potential synergies and tradeoffs among individual reproductive components may play a much larger role in influencing reproductive success over a male’s lifetime than within a single breeding season. In particular, positive within-pair/extra-pair covariances in the opportunity for selection, as observed here, indicate that there are large potential lifetime fitness benefits for males able to capitalize on mating with extra-pair females while still investing in behaviors that enhance *W*_*RS*_ with their socially-paired mate(s). Previous research from this and other systems indicate that local breeding synchrony likely plays a role in the ability of males to copulate with extra-pair female neighbours while maintaining their own within-pair reproductive success (Stutchbury and Neudorf 1998; Chuang et al. 1999; Webster et al. 2001; Stewart et al. 2010). By mating with synchronously fertile females on nearby adjacent territories, males of some species can reduce the risk of losing within-pair paternity of their own offspring via reduced mate guarding while seeking potential extra-pair mates over greater distances throughout the population (Kempenaers 1997; Weatherhead 1997; Reitsma et al. 2018). In our system, males on food-rich territories with double-brooding females may have increased access to neighboring females that likewise double-brood given greater food resources (Holmes et al. 1996; Nagy and Holmes 2005b), increasing the overall availability of both within-pair and extra-pair offspring for the focal male to sire. Future work investigating environmental drivers of the opportunity for selection should thus consider the role of local breeding synchrony as it relates to positive covariances between the *W*_*RS*_ and *E*_*RS*_ pathways.

Unlike our analysis of annual reproductive success, male lifetime reproductive success was more strongly influenced by female breeding quality and paternity allocation than by the number of mates a male acquired. Of the components of (co)variance found to have a substantial influence on the opportunity for sexual selection in lifetime reproductive success, six had significant, positive effects on 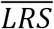 in the population (Figure 5). Surprisingly, neither the number of within-pair nor number of extra-pair mates had any significant effect on 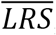, contrary to our results for 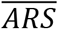 (Figure 4). Instead, all individual components of variance related to female fecundity (*B*_*w*_, *F*_*w*_, *B*_*e*_, *F*_*e*_) and male siring success (*P*_*w*_, *P*_*e*_) had significant positive effects on 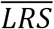. Further, the interaction between number of within-pair mates and their fecundity per brood, as well as the interaction between number of extra-pair mates and fecundity per brood of those mates were found to have significant negative effects on 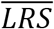 (Figure 5), indicating that male black-throated blue warblers experience a lifetime trade-off in terms of the number and the quality of their mates through both the *W*_*RS*_ and *E*_*RS*_ pathways. These results are consistent with the hypothesis that males with multiple within-pair and/or extra-pair mates suffer reproductive loses annually compared to monogamous males (e.g. Dunn and Robertson 1993; Poirier et al. 2004; Reitsma et al. 2018). However, such patterns of negative covariance and lower reproductive success are not widely reported over the lifetimes of breeding males, in particular when contrasted with the apparent positive effects of more within-pair and extra-pair mates within a single breeding season. Taken together, our results suggest that males in this population may sacrifice short-term reproductive gains (i.e. higher 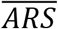 via more within-pair and/or extra-pair mates) for longer-term fitness benefits (represented by higher 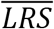).

Standardized variance, commonly denoted *I*_s_ and which traditionally represents the metric of comparing the opportunity for selection among studies, was moderate for male annual reproductive success in our population, but relatively high for lifetime reproductive success (Table 1, Table 2). This is reflected by the greater degree of reproductive skew among males in terms of lifetime reproduction (Figure 2), where certain males in our population achieve exceptionally high output of within-pair and/or extra-pair offspring. Standardized variance in *T*_*ARS*_ was comparable to other species where within-pair success appears to play a larger role than extra-pair success in the opportunity for selection (e.g. savannah sparrows – Freeman-Gallant et al. 2005; roe deer – Vanpé et al. 2008), and considerably lower than among species where polygynous (i.e. extra-pair) mating drives the opportunity for selection in annual reproduction (e.g. fruit bats – Storz et al. 2001; sand gobies – Jones et al. 2001). Likewise, standardized variance in *T*_*LRS*_ was again lower than comparable studies where extra-pair reproduction drives variance in male fitness over their lifetimes (e.g. red-backed fairy wrens – Webster et al. 2007; song sparrows – Lebigre et al. 2015), but higher than estimates of standardized variance among primarily monogamous populations (e.g. California mouse – Ribble 1992; prairie voles – Shuster et al. 2019). Thus, while overall variance in male reproductive success in our population may be less skewed than observed in systems with higher rates of extra-pair paternity, our results indicate that the potential fitness benefits of males allocating more resources to securing *W*_*RS*_, in tandem with opportunistically seeking *E*_*RS*_, leads to a greater overall opportunity for selection than expected if male time/energy were directed to their within-pair mates alone.

Overall, our results suggest that variance in within-pair reproductive success clearly drives the opportunity for selection in this multi-brooded population. Our ability to quantify annual variation in the effects of *W*_*RS*_ and *E*_*RS*_ as well as compare drivers of variance in *T*_*ARS*_ and *T*_*LRS*_ using this long-term dataset provides unique insight into how the opportunity for selection operates annually and over the lifetimes of males in a multi-brooded, socially monogamous but genetically polygynous species. While the components of (co)variance which most strongly affect both the opportunity for selection and realized reproductive success differed somewhat between short (i.e. annual) and longer (i.e. lifetime) timescales, components of *W*_*RS*_ consistently accounted for more total variance in *T*_*ARS*_ and *T*_*LRS*_, and had greater contributions to higher mean annual and lifetime reproductive success, than did components of *E*_*RS*_. The potential for annual reproductive tradeoffs between within-pair and extra-pair components of variance were outweighed by the overall positive covariance between *W*_*RS*_ and *E*_*RS*_. Thus, males able to maximize *W*_*RS*_ likely achieve higher overall lifetime fitness via both the within-pair and extra-pair reproductive pathways. This finding differs from comparable studies where greater extra-pair reproductive success among males is either uncorrelated with or can come at the expense of within-pair success (i.e. minimal or negative covariance; Webster et al. 1995; Freeman-Gallant et al. 2005; Webster et al. 2007, Lebigre et al. 2012; Taff et al. 2013), and thus hints at a potential ‘winner take all’ style of mating system within this socially monogamous yet genetically polygynous species. Minimal observed effects of annual variation in socio-environmental factors on the influences of components of *W*_*RS*_ indicate that consistent selection pressures may offer some males (e.g. those on food-rich territories, Kaiser et al. 2015, 2017) the opportunity to capitalize on long-term reproductive benefits (e.g. double-brooding by their socially-paired female), even if such benefits result in short term losses in annual reproductive success via reduced effort in seeking additional mates. In contrast, males on food-poor territories may experience more selective pressure to allocate resources towards extra-pair reproduction given their lower probability of within-pair success (Kaiser et al. 2015, 2017), leading to greater variance in components of *E*_*RS*_ in this subset of males. While our current study does not directly determine which phenotypic or ecological traits are influenced by the opportunity for selection (e.g. secondary sexual signals, competitive ability for territories), future studies should seek to determine if and how the opportunity for selection differs between high vs. low-quality territories in this and other systems, particularly as changing climate conditions alter breeding-season length and potentially reduce population-wide variance in key components of reproduction such as the ability of females to initiate multiple broods. Such studies, especially with regard to variance in reproduction among females, will not only provide unique insight into how habitat type can influence selection and the lifetime fitness, but also advance our understanding of how changing socio-environmental conditions can ultimately alter the mating systems of wild populations in seasonal environments.

## Author contributions

R.R.G. and M.S.W. conceived the study, and R.R.G. conducted statistical analyses and wrote the manuscript with input from all authors. M.T.H, S.A.K, T.S.S. and M.S.W. collected data, and S.A.K. conducted parentage assignment.

## Acknowledgements

We thank the many undergraduates and field technicians that contributed towards data collection, G.J. Colbeck, E.R.A. Cramer, and S. Bartos-Smith for assistance in the laboratory and with genotyping, and R.T. Holmes, M.P. Ayers, and N.L. Rodenhouse for logistic support. This research is a contribution of the Hubbard Brook Ecosystem Study, part of the Long-Term Ecological Research network supported by the National Science Foundation (grants awarded to Cornell University [0640470], the Smithsonian Institution [0640732] and Wellesley College [064082300]). R.R.G was supported by a postdoctoral grant from the Natural Sciences and Engineering Research Council of Canada.

## Data Accessibility

All data underlying these analyses will be uploaded to Dryad upon acceptance

## Additional Files

Supporting Information: Supporting Information sections S1-S3

## Notes

### Competing Interest Statement

The authors have declared no competing interest.

