## Supporting Information for "Variance in within-pair reproductive success influences the opportunity for selection annually and over the lifetimes of males in a multi-brooded songbird"

### Supporting Information S1 – Details of variance term calculations

Variance partitioning of male total reproductive success ($T_{RS}$) follows the general formula:

$$Var(T_{RS}) = Var(W_{RS}) + Var(E_{RS}) + 2\ Cov(W_{RS}, E_{RS}),$$

where *Var* represents the variance in each component and *Cov(W,E)* the covariance between within-pair and extra-pair reproductive success (Webster et al. 1995). This expression can be further decomposed to each component source of within-pair and extra-pair reproductive success (Figure 1) by accounting for mean population-wide values for the remaining terms, following Bohrnstedt and Goldberger (1969). For instance, calculating variance in the number of within-pair mates ($M_w$) requires multiplying the squared mean number of broods per WP mate by the squared means of fecundity per brood per WP mate and the proportion of WP offspring sired per brood: $\bar{B}_w^2\bar{F}_w^2\bar{P}_w^2Var(M_w)$. Likewise, calculating covariance terms for within-pair or extra-pair success requires multiplying covariance by 2× the squared mean value of the remaining term(s), times the mean value of the terms of interest. For instance, the covariance between number of within-pair mates ($M_w$) and number of broods per within-pair mate ($B_w$) is calculated as: $2\bar{M}_w\bar{B}_w\bar{F}_w^2\bar{P}_w^2Cov(M_w, B_w)$. However, when calculated covariance between within-pair and extra-pair terms, we forgo squaring any mean values and simply calculate covariance as 2× the mean value for each remaining term. For instance, covariance between the number of within-pair and extra-pair mates $Cov(M_w, M_e)$ is calculated as: $2\bar{B}_w\bar{F}_w\bar{P}_w\bar{B}_e\bar{F}_e\bar{P}_eCov(M_w, M_e)$. Table S1 provides a full list of formulas used to calculate each variance component, as well as the simplified term used in-text for ease of interpretation. See Bohrnstedt and Goldberger (1969) and Webster et al. (1995) for further details and equation proofs.

**Table S1:** Terms used to describe within-pair (WP) and extra-pair (EP) variance components in-text, their full-expanded formula, and biological interpretation.

| Source | Term used in text | Formula | Interpretation |
|---|---|---|---|
| Within-pair terms | $M_w$ | $\bar{B}_w^2\bar{F}_w^2\bar{P}_w^2Var(M_w)$ | Number of WP mates |
| | $B_w$ | $\bar{M}_w^2\bar{F}_w^2\bar{P}_w^2Var(B_w)$ | Number of broods per WP mate |
| | $F_w$ | $\bar{M}_w^2\bar{B}_w^2\bar{P}_w^2Var(F_w)$ | Fecundity per brood per WP mate |
| | $P_w$ | $\bar{M}_w^2\bar{B}_w^2\bar{F}_w^2Var(P_w)$ | Proportion of young sired per brood per WP mate |
| | $Cov(M_w,B_w)$ | $2\bar{M}_w\bar{B}_w\bar{F}_w^2\bar{P}_w^2Cov(M_w,B_w)$ | Covariance – WP mates, broods per WP mate |
| | $Cov(M_w,F_w)$ | $2\bar{M}_w\bar{B}_w^2\bar{F}_w\bar{P}_w^2Cov(M_w,F_w)$ | Covariance – WP mates, fecundity per brood per WP mate |
| | $Cov(M_w,P_w)$ | $2\bar{M}_w\bar{B}_w^2\bar{F}_w^2\bar{P}_wCov(M_w,P_w)$ | Covariance – WP mates, proportion sired per brood per WP mate |
| | $Cov(B_w,F_w)$ | $2\bar{M}_w^2\bar{B}_w\bar{F}_w\bar{P}_w^2Cov(B_w,F_w)$ | Covariance – broods per WP mate, fecundity per brood per WP mate |
| | $Cov(B_w,P_w)$ | $2\bar{M}_w^2\bar{B}_w\bar{F}_w^2\bar{P}_wCov(B_w,P_w)$ | Covariance – broods per WP mate, proportion sired per brood per WP mate |
| | $Cov(F_e,P_e)$ | $2\bar{M}_w^2\bar{B}_w^2\bar{F}_w\bar{P}_wCov(F_e,P_e)$ | Covariance – fecundity per brood per WP mate, proportion sired per brood per WP mate |
| Extra-pair terms | $M_e$ | $\bar{B}_e^2\bar{F}_e^2\bar{P}_e^2Var(M_e)$ | Number of EP mates |
| | $B_e$ | $\bar{M}_e^2\bar{F}_e^2\bar{P}_e^2Var(B_e)$ | Number of broods per EP mate |
| | $F_e$ | $\bar{M}_e^2\bar{B}_e^2\bar{P}_e^2Var(F_e)$ | Fecundity per brood per EP mate |

| Source | Term used in text | Formula | Interpretation |
|---|---|---|---|
| | $P_e$ | $\bar{M}_e^2\bar{B}_e^2\bar{F}_e^2Var(P_e)$ | Proportion of young sired per brood per EP mate |
| | $Cov(M_e, B_e)$ | $2\bar{M}_e\bar{B}_e\bar{F}_e^2\bar{P}_e^2Cov(M_e, B_e)$ | Covariance – EP mates, broods per EP mate |
| | $Cov(M_e, F_e)$ | $2\bar{M}_e\bar{B}_e^2\bar{F}_e\bar{P}_e^2Cov(M_e, F_e)$ | Covariance – EP mates, fecundity per brood per EP mate |
| | $Cov(M_e, P_e)$ | $2\bar{M}_e\bar{B}_e^2\bar{F}_e^2\bar{P}_eCov(M_e, P_e)$ | Covariance – EP mates, proportion sired per brood per EP mate |
| | $Cov(B_e, F_e)$ | $2\bar{M}_e^2\bar{B}_e\bar{F}_e\bar{P}_e^2Cov(B_e, F_e)$ | Covariance – broods per EP mate, fecundity per brood per EP mate |
| | $Cov(B_e, P_e)$ | $2\bar{M}_e^2\bar{B}_e\bar{F}_e^2\bar{P}_eCov(B_e, P_e)$ | Covariance – broods per EP mate, proportion sired per brood per EP mate |
| | $Cov(F_e, P_e)$ | $2\bar{M}_e^2\bar{B}_e^2\bar{F}_e\bar{P}_eCov(F_e, P_e)$ | Covariance – fecundity per brood per EP mate, proportion sired per brood per EP mate |
| Within-pair and extra-pair covariances | $Cov(M_w, M_e)$ | $2\bar{B}_w\bar{F}_w\bar{P}_w\bar{B}_e\bar{F}_e\bar{P}_eCov(M_w, M_e)$ | Covariance – WP mates, EP mates |
| | $Cov(M_w, B_e)$ | $2\bar{B}_w\bar{F}_w\bar{P}_w\bar{M}_e\bar{F}_e\bar{P}_eCov(M_w, B_e)$ | Covariance – WP mates, broods per EP mate |
| | $Cov(M_w, F_e)$ | $2\bar{B}_w\bar{F}_w\bar{P}_w\bar{M}_e\bar{B}_e\bar{P}_eCov(M_w, F_e)$ | Covariance – WP mates, fecundity per brood per EP mate |
| | $Cov(M_w, P_e)$ | $2\bar{B}_w\bar{F}_w\bar{P}_w\bar{M}_e\bar{B}_e\bar{F}_eCov(M_w, P_e)$ | Covariance – WP mates, proportion sired per brood per EP mate |
| | $Cov(B_w, M_e)$ | $2\bar{M}_w\bar{F}_w\bar{P}_w\bar{B}_e\bar{F}_e\bar{P}_eCov(B_w, M_e)$ | Covariance – broods per WP mate, EP mates |
| | $Cov(B_w, B_e)$ | $2\bar{M}_w\bar{F}_w\bar{P}_w\bar{M}_e\bar{F}_e\bar{P}_eCov(B_w, B_e)$ | Covariance – broods per WP mate, broods per EP mate |

| Source | Term used in text | Formula | Interpretation |
|---|---|---|---|
| | $Cov(B_w, F_e)$ | $2\bar{M}_w\bar{F}_w\bar{P}_w\bar{M}_e\bar{B}_e\bar{P}_eCov(B_w, F_e)$ | Covariance – broods per WP mate, fecundity per brood per EP mate |
| | $Cov(B_w, P_e)$ | $2\bar{M}_w\bar{F}_w\bar{P}_w\bar{M}_e\bar{B}_e\bar{F}_eCov(B_w, P_e)$ | Covariance – broods per WP mate, proportion sired per brood per EP mate |
| | $Cov(F_w, M_e)$ | $2\bar{M}_w\bar{B}_w\bar{P}_w\bar{B}_e\bar{F}_e\bar{P}_eCov(F_w, M_e)$ | Covariance – fecundity per brood per WP mate, EP mates |
| | $Cov(F_w, B_e)$ | $2\bar{M}_w\bar{B}_w\bar{P}_w\bar{M}_e\bar{F}_e\bar{P}_eCov(F_w, B_e)$ | Covariance – fecundity per brood per WP mate, broods per EP mate |
| | $Cov(F_w, F_e)$ | $2\bar{M}_w\bar{B}_w\bar{P}_w\bar{M}_e\bar{B}_e\bar{P}_eCov(F_w, F_e)$ | Covariance – fecundity per brood per WP mate, fecundity per brood per EP mate |
| | $Cov(F_w, P_e)$ | $2\bar{M}_w\bar{B}_w\bar{P}_w\bar{M}_e\bar{B}_e\bar{F}_eCov(F_w, P_e)$ | Covariance – fecundity per brood per WP mate, proportion sired per brood per EP mate |
| | $Cov(P_w, M_e)$ | $2\bar{M}_w\bar{B}_w\bar{F}_w\bar{B}_e\bar{F}_e\bar{P}_eCov(P_w, M_e)$ | Covariance – proportion sired per brood per WP mate, EP mates |
| | $Cov(P_w, B_e)$ | $2\bar{M}_w\bar{B}_w\bar{F}_w\bar{M}_e\bar{F}_e\bar{P}_eCov(P_w, B_e)$ | Covariance – proportion sired per brood per WP mate, broods per EP mate |
| | $Cov(P_w, F_e)$ | $2\bar{M}_w\bar{B}_w\bar{F}_w\bar{M}_e\bar{B}_e\bar{P}_eCov(P_w, F_e)$ | Covariance – proportion sired per brood per WP mate, fecundity per brood per EP mate |
| | $Cov(P_w, P_e)$ | $2\bar{M}_w\bar{B}_w\bar{F}_w\bar{M}_e\bar{B}_e\bar{F}_eCov(P_w, P_e)$ | Covariance – proportion sired per brood per WP mate, proportion sired per brood per EP mate |

**Supporting Information S2 – Full output from ‘top models subset’ of annual reproductive success**

**Table S2:** Parameter estimates for all six models in ‘top models subset’ (ΔAICc ≤ 7 from top model, arranged by increasing ΔAICc) describing the effects of key variance components on mean annual reproductive success in our population of black-throated blue warblers. ‘Weight’ refers to the AIC weight of the focal model. Blank cells indicate where a term was not included in a particular model.

| | Intercept | $M_w$ | $B_w$ | $P_w$ | $M_e$ | $B_e$ | $B_w$ x $B_e$ | *df* | ΔAICc | Weight |
|---|---|---|---|---|---|---|---|---|---|---|
| 1 | 1.42 | 0.06 | 0.15 | 0.33 | 0.21 | | | 7 | 0 | 0.29 |
| 2 | 1.42 | 0.06 | 0.14 | 0.33 | 0.21 | 0.04 | | 8 | 0.33 | 0.25 |
| 3 | 1.42 | 0.06 | 0.13 | 0.33 | 0.21 | 0.04 | 0.04 | 9 | 1.06 | 0.17 |
| 4 | 1.43 | | 0.15 | 0.33 | 0.21 | | | 6 | 1.65 | 0.13 |
| 5 | 1.43 | | 0.14 | 0.32 | 0.21 | 0.04 | | 7 | 2.12 | 0.10 |
| 6 | 1.42 | | 0.13 | 0.32 | 0.21 | 0.04 | 0.03 | 8 | 3.22 | 0.06 |

**Table S3:** Parameter estimates, 95% confidence intervals (CI), and z-values for final averaged model describing the effects of key variance components on mean annual reproductive success.

| Component | Estimate | Lower CI | Upper CI | z value |
|---|---|---|---|---|
| $M_w$ | 0.04 | 0.003 | 0.11 | 1.16 |
| $B_w$ | 0.14 | 0.09 | 0.20 | 4.84 |
| $P_w$ | 0.33 | 0.26 | 0.39 | 9.73 |
| $M_e$ | 0.21 | 0.17 | 0.26 | 9.07 |
| $B_e$ | 0.02 | -0.02 | 0.10 | 0.75 |
| $B_w$ x $B_e$ | 0.01 | -0.03 | 0.09 | 0.39 |

**Supporting Information S3 – Full output from ‘top models subset’ of lifetime reproductive success**

**Table S4:** Parameter estimates for all 147 models in ‘top models subset’ (ΔAICc ≤ 7 from top model, arranged by increasing ΔAICc) describing the effects of key variance components on mean lifetime reproductive success in our population of black-throated blue warblers. ‘Weight’ refers to the AIC weight of the focal model. Blank cells indicate where a term was not included in a particular model.

| | Intercept | $M_w$ | $B_w$ | $F_w$ | $P_w$ | $M_w$ x $F_w$ | $M_e$ | $B_e$ | $F_e$ | $P_e$ | $M_e$ x $F_e$ | $M_w$ x $M_e$ | $M_w$ x $F_e$ | $M_w$ x $P_e$ | $B_w$ x $B_e$ | $B_w$ x $P_e$ | $F_w$ x $M_e$ | $F_w$ x $F_e$ | df | ΔAICc | Weight |
|---|---|---|---|---|---|---|---|---|---|---|---|---|---|---|---|---|---|---|---|---|---|
| 1 | 1.91 | 0.07 | 0.20 | 0.34 | 0.29 | -0.04 | 0.04 | 0.13 | 0.33 | 0.18 | -0.04 | | | | | | | -0.05 | 14 | 0.00 | 0.04 |
| 2 | 1.92 | 0.09 | 0.21 | 0.32 | 0.28 | -0.06 | -0.03 | 0.12 | 0.40 | 0.18 | -0.04 | | | | | | 0.10 | -0.14 | 15 | 0.18 | 0.04 |
| 3 | 1.91 | 0.09 | 0.21 | 0.33 | 0.29 | -0.06 | 0.03 | 0.13 | 0.32 | 0.18 | -0.05 | | | | | | | | 13 | 0.85 | 0.03 |
| 4 | 1.91 | 0.07 | 0.20 | 0.34 | 0.28 | -0.04 | 0.03 | 0.13 | 0.34 | 0.19 | -0.04 | | | -0.03 | | | | -0.05 | 15 | 0.91 | 0.03 |
| 5 | 1.92 | 0.10 | 0.21 | 0.33 | 0.28 | -0.06 | -0.04 | 0.12 | 0.41 | 0.19 | -0.04 | | | -0.03 | | | 0.10 | -0.15 | 16 | 1.12 | 0.03 |
| 6 | 1.92 | 0.12 | 0.21 | 0.31 | 0.28 | -0.07 | -0.05 | 0.12 | 0.40 | 0.19 | -0.05 | 0.14 | -0.16 | | | | | | 15 | 1.15 | 0.02 |
| 7 | 1.92 | 0.09 | 0.21 | 0.34 | 0.28 | -0.06 | -0.03 | 0.12 | 0.40 | 0.19 | -0.04 | 0.20 | -0.18 | | | | -0.06 | | 16 | 1.28 | 0.02 |
| 8 | 1.92 | 0.09 | 0.21 | 0.33 | 0.28 | -0.06 | -0.04 | 0.12 | 0.41 | 0.19 | -0.04 | 0.15 | -0.13 | | | | | -0.05 | 16 | 1.34 | 0.02 |
| 9 | 1.89 | | 0.19 | 0.34 | 0.28 | | 0.02 | 0.13 | 0.38 | 0.18 | -0.04 | | | | | | | -0.08 | 12 | 1.35 | 0.02 |
| 10 | 1.91 | 0.06 | 0.20 | 0.35 | 0.28 | -0.05 | 0.03 | 0.13 | 0.34 | 0.18 | -0.05 | 0.03 | | | | | | -0.07 | 15 | 1.52 | 0.02 |
| 11 | 1.91 | 0.10 | 0.21 | 0.31 | 0.29 | -0.05 | 0.03 | 0.13 | 0.33 | 0.18 | -0.04 | | -0.03 | | | | | | 14 | 1.88 | 0.02 |
| 12 | 1.91 | 0.07 | 0.21 | 0.34 | 0.29 | -0.05 | 0.06 | 0.13 | 0.31 | 0.18 | -0.05 | | | | | | -0.03 | | 14 | 1.89 | 0.02 |

| | Intercept | $M_w$ | $B_w$ | $F_w$ | $P_w$ | $M_w$ x $F_w$ | $M_e$ | $B_e$ | $F_e$ | $P_e$ | $M_e$ x $F_e$ | $M_w$ x $M_e$ | $M_w$ x $F_e$ | $M_w$ x $P_e$ | $B_w$ x $B_e$ | $B_w$ x $P_e$ | $F_w$ x $M_e$ | $F_w$ x $F_e$ | df | ΔAICc | Weight |
|---|---|---|---|---|---|---|---|---|---|---|---|---|---|---|---|---|---|---|---|---|---|
| 13 | 1.91 | 0.07 | 0.20 | 0.33 | 0.28 | -0.04 | 0.04 | 0.13 | 0.33 | 0.18 | -0.04 | | | | | -0.01 | | -0.05 | 15 | 2.21 | 0.01 |
| 14 | 1.91 | 0.07 | 0.20 | 0.34 | 0.29 | -0.04 | 0.04 | 0.13 | 0.33 | 0.18 | -0.04 | | | | 0.00 | | | -0.05 | 15 | 2.25 | 0.01 |
| 15 | 1.91 | 0.07 | 0.20 | 0.34 | 0.29 | -0.04 | 0.04 | 0.13 | 0.33 | 0.18 | -0.04 | | 0.01 | | | | | -0.05 | 15 | 2.25 | 0.01 |
| 16 | 1.92 | 0.13 | 0.21 | 0.31 | 0.28 | -0.07 | -0.06 | 0.12 | 0.41 | 0.19 | -0.04 | 0.14 | -0.16 | -0.03 | | | | | 16 | 2.26 | 0.01 |
| 17 | 1.92 | 0.09 | 0.21 | 0.33 | 0.28 | -0.06 | -0.03 | 0.12 | 0.40 | 0.19 | -0.05 | 0.01 | | | | | 0.09 | -0.14 | 16 | 2.30 | 0.01 |
| 18 | 1.92 | 0.09 | 0.21 | 0.32 | 0.28 | -0.06 | -0.03 | 0.12 | 0.40 | 0.18 | -0.04 | | | | -0.01 | | 0.11 | -0.14 | 16 | 2.36 | 0.01 |
| 19 | 1.92 | 0.09 | 0.21 | 0.33 | 0.28 | -0.06 | -0.03 | 0.12 | 0.40 | 0.19 | -0.05 | | 0.01 | | | | 0.10 | -0.15 | 16 | 2.43 | 0.01 |
| 20 | 1.92 | 0.10 | 0.21 | 0.32 | 0.28 | -0.06 | -0.03 | 0.12 | 0.40 | 0.18 | -0.04 | | | | | 0.00 | 0.10 | -0.14 | 16 | 2.45 | 0.01 |
| 21 | 1.92 | 0.10 | 0.21 | 0.33 | 0.28 | -0.06 | -0.05 | 0.12 | 0.41 | 0.19 | -0.04 | 0.14 | -0.14 | -0.03 | | | | -0.05 | 17 | 2.54 | 0.01 |
| 22 | 1.92 | 0.10 | 0.21 | 0.33 | 0.28 | -0.06 | -0.04 | 0.12 | 0.40 | 0.19 | -0.04 | 0.20 | -0.19 | -0.02 | | | -0.05 | | 17 | 2.55 | 0.01 |
| 23 | 1.91 | 0.09 | 0.21 | 0.33 | 0.29 | -0.06 | 0.02 | 0.13 | 0.33 | 0.19 | -0.05 | | | -0.02 | | | | | 14 | 2.55 | 0.01 |
| 24 | 1.91 | 0.10 | 0.21 | 0.31 | 0.29 | -0.05 | 0.02 | 0.13 | 0.34 | 0.19 | -0.04 | | -0.04 | -0.03 | | | | | 15 | 2.74 | 0.01 |
| 25 | 1.91 | 0.09 | 0.21 | 0.32 | 0.29 | -0.06 | 0.04 | 0.13 | 0.32 | 0.18 | -0.05 | -0.01 | | | | | | | 14 | 2.83 | 0.01 |
| 26 | 1.91 | 0.06 | 0.20 | 0.35 | 0.28 | -0.05 | 0.02 | 0.13 | 0.35 | 0.19 | -0.04 | 0.02 | | -0.02 | | | | -0.07 | 16 | 2.87 | 0.01 |
| 27 | 1.88 | | 0.19 | 0.35 | 0.28 | | 0.07 | 0.14 | 0.33 | 0.18 | -0.04 | | | | | | -0.07 | | 12 | 2.95 | 0.01 |
| 28 | 1.92 | 0.09 | 0.21 | 0.33 | 0.29 | -0.06 | 0.03 | 0.13 | 0.32 | 0.18 | -0.05 | | | | -0.01 | | | | 14 | 3.07 | 0.01 |
| 29 | 1.91 | 0.09 | 0.21 | 0.33 | 0.29 | -0.06 | 0.03 | 0.13 | 0.32 | 0.18 | -0.05 | | | | | 0.00 | | | 14 | 3.08 | 0.01 |
| 30 | 1.91 | 0.07 | 0.20 | 0.34 | 0.28 | -0.04 | 0.03 | 0.13 | 0.34 | 0.19 | -0.04 | | -0.01 | -0.03 | | | | -0.05 | 16 | 3.18 | 0.01 |

| | Intercept | $M_w$ | $B_w$ | $F_w$ | $P_w$ | $M_w$ x $F_w$ | $M_e$ | $B_e$ | $F_e$ | $P_e$ | $M_e$ x $F_e$ | $M_w$ x $M_e$ | $M_w$ x $F_e$ | $M_w$ x $P_e$ | $B_w$ x $B_e$ | $B_w$ x $P_e$ | $F_w$ x $M_e$ | $F_w$ x $F_e$ | df | ΔAICc | Weight |
|---|---|---|---|---|---|---|---|---|---|---|---|---|---|---|---|---|---|---|---|---|---|
| 31 | 1.91 | 0.07 | 0.20 | 0.34 | 0.28 | -0.04 | 0.03 | 0.13 | 0.34 | 0.19 | -0.04 | | | -0.03 | | 0.00 | | -0.05 | 16 | 3.18 | 0.01 |
| 32 | 1.91 | 0.07 | 0.21 | 0.34 | 0.29 | -0.05 | 0.05 | 0.13 | 0.31 | 0.19 | -0.04 | | | -0.02 | | | -0.03 | | 15 | 3.18 | 0.01 |
| 33 | 1.91 | 0.07 | 0.20 | 0.34 | 0.28 | -0.04 | 0.03 | 0.13 | 0.34 | 0.19 | -0.04 | | | -0.03 | 0.00 | | | -0.05 | 16 | 3.19 | 0.01 |
| 34 | 1.89 | 0.03 | 0.19 | 0.32 | 0.28 | | 0.02 | 0.13 | 0.37 | 0.18 | -0.04 | | | | | | | -0.08 | 13 | 3.25 | 0.01 |
| 35 | 1.92 | 0.12 | 0.21 | 0.31 | 0.28 | -0.07 | -0.05 | 0.12 | 0.41 | 0.19 | -0.05 | 0.14 | -0.16 | | -0.01 | | | | 16 | 3.29 | 0.01 |
| 36 | 1.92 | 0.10 | 0.21 | 0.33 | 0.28 | -0.06 | -0.05 | 0.12 | 0.41 | 0.19 | -0.04 | | | -0.03 | -0.01 | | 0.10 | -0.15 | 17 | 3.35 | 0.01 |
| 37 | 1.92 | 0.12 | 0.21 | 0.30 | 0.28 | -0.07 | -0.05 | 0.12 | 0.40 | 0.19 | -0.05 | 0.14 | -0.16 | | | -0.01 | | | 16 | 3.38 | 0.01 |
| 38 | 1.92 | 0.09 | 0.21 | 0.33 | 0.28 | -0.06 | -0.04 | 0.12 | 0.41 | 0.19 | -0.04 | 0.00 | | -0.03 | | | 0.10 | -0.15 | 17 | 3.43 | 0.01 |
| 39 | 1.92 | 0.10 | 0.21 | 0.32 | 0.28 | -0.06 | -0.05 | 0.12 | 0.41 | 0.19 | -0.04 | | 0.00 | -0.03 | | | 0.10 | -0.14 | 17 | 3.43 | 0.01 |
| 40 | 1.92 | 0.10 | 0.21 | 0.33 | 0.28 | -0.06 | -0.04 | 0.12 | 0.41 | 0.19 | -0.04 | | | -0.03 | | 0.00 | 0.10 | -0.15 | 17 | 3.43 | 0.01 |
| 41 | 1.92 | 0.09 | 0.21 | 0.34 | 0.28 | -0.06 | -0.03 | 0.12 | 0.40 | 0.19 | -0.04 | 0.20 | -0.19 | | -0.01 | | -0.06 | | 17 | 3.50 | 0.01 |
| 42 | 1.89 | | 0.19 | 0.34 | 0.28 | | 0.02 | 0.13 | 0.38 | 0.18 | -0.04 | | | | -0.01 | | | -0.08 | 13 | 3.53 | 0.01 |
| 43 | 1.88 | | 0.19 | 0.34 | 0.28 | | 0.02 | 0.13 | 0.38 | 0.18 | -0.04 | | | | | -0.01 | | -0.08 | 13 | 3.54 | 0.01 |
| 44 | 1.92 | 0.09 | 0.21 | 0.33 | 0.28 | -0.06 | -0.04 | 0.12 | 0.41 | 0.19 | -0.04 | 0.15 | -0.13 | | -0.01 | | | -0.05 | 17 | 3.54 | 0.01 |
| 45 | 1.92 | 0.09 | 0.21 | 0.33 | 0.28 | -0.06 | -0.03 | 0.12 | 0.40 | 0.19 | -0.04 | 0.20 | -0.18 | | | -0.01 | -0.06 | | 17 | 3.56 | 0.01 |
| 46 | 1.89 | | 0.19 | 0.34 | 0.28 | | 0.01 | 0.13 | 0.38 | 0.18 | -0.04 | | | | | | 0.00 | -0.08 | 13 | 3.58 | 0.01 |
| 47 | 1.92 | 0.09 | 0.21 | 0.34 | 0.28 | -0.06 | -0.03 | 0.12 | 0.40 | 0.19 | -0.04 | 0.19 | -0.17 | | | | -0.04 | -0.02 | 17 | 3.58 | 0.01 |
| 48 | 1.92 | 0.09 | 0.21 | 0.33 | 0.28 | -0.06 | -0.04 | 0.12 | 0.41 | 0.19 | -0.04 | 0.15 | -0.13 | | | 0.00 | | -0.05 | 17 | 3.63 | 0.01 |

| | Intercept | $M_w$ | $B_w$ | $F_w$ | $P_w$ | $M_w$ x $F_w$ | $M_e$ | $B_e$ | $F_e$ | $P_e$ | $M_e$ x $F_e$ | $M_w$ x $M_e$ | $M_w$ x $F_e$ | $M_w$ x $P_e$ | $B_w$ x $B_e$ | $B_w$ x $P_e$ | $F_w$ x $M_e$ | $F_w$ x $F_e$ | df | ΔAICc | Weight |
|---|---|---|---|---|---|---|---|---|---|---|---|---|---|---|---|---|---|---|---|---|---|
| 49 | 1.89 | 0.03 | 0.19 | 0.32 | 0.28 | | 0.01 | 0.13 | 0.38 | 0.19 | -0.03 | | | -0.03 | | | | -0.09 | 14 | 3.77 | 0.01 |
| 50 | 1.92 | 0.06 | 0.20 | 0.35 | 0.28 | -0.05 | 0.03 | 0.13 | 0.34 | 0.18 | -0.05 | 0.03 | | | 0.00 | | | -0.07 | 16 | 3.79 | 0.01 |
| 51 | 1.91 | 0.06 | 0.20 | 0.35 | 0.28 | -0.05 | 0.03 | 0.13 | 0.34 | 0.18 | -0.05 | 0.03 | | | | 0.00 | | -0.07 | 16 | 3.80 | 0.01 |
| 52 | 1.91 | 0.08 | 0.21 | 0.33 | 0.29 | -0.05 | 0.05 | 0.13 | 0.32 | 0.18 | -0.04 | | -0.02 | | | | -0.02 | | 15 | 3.88 | 0.01 |
| 53 | 1.87 | | 0.19 | 0.37 | 0.29 | | | 0.13 | 0.33 | 0.17 | | | | | | | | -0.10 | 10 | 4.04 | 0.01 |
| 54 | 1.91 | 0.06 | 0.21 | 0.35 | 0.29 | -0.05 | 0.06 | 0.13 | 0.31 | 0.18 | -0.05 | 0.01 | | | | | -0.04 | | 15 | 4.04 | 0.01 |
| 55 | 1.91 | 0.10 | 0.21 | 0.31 | 0.29 | -0.05 | 0.03 | 0.12 | 0.33 | 0.18 | -0.04 | | -0.03 | | | -0.01 | | | 15 | 4.06 | 0.01 |
| 56 | 1.91 | 0.10 | 0.21 | 0.32 | 0.29 | -0.05 | 0.03 | 0.13 | 0.33 | 0.18 | -0.04 | | -0.03 | | -0.01 | | | | 15 | 4.10 | 0.01 |
| 57 | 1.91 | 0.07 | 0.21 | 0.34 | 0.29 | -0.05 | 0.06 | 0.13 | 0.31 | 0.18 | -0.05 | | | | | -0.01 | -0.03 | | 15 | 4.11 | 0.01 |
| 58 | 1.91 | 0.09 | 0.21 | 0.32 | 0.29 | -0.05 | 0.03 | 0.13 | 0.33 | 0.19 | -0.04 | -0.02 | | -0.02 | | | | | 15 | 4.14 | 0.01 |
| 59 | 1.91 | 0.07 | 0.21 | 0.34 | 0.29 | -0.05 | 0.06 | 0.13 | 0.31 | 0.18 | -0.05 | | | | 0.00 | | -0.03 | | 15 | 4.15 | 0.01 |
| 60 | 1.88 | 0.05 | 0.20 | 0.37 | 0.29 | -0.03 | | 0.13 | 0.30 | 0.18 | | | | -0.04 | | | | -0.08 | 13 | 4.18 | 0.01 |
| 61 | 1.91 | 0.07 | 0.20 | 0.33 | 0.28 | -0.04 | 0.04 | 0.13 | 0.33 | 0.18 | -0.04 | | | | -0.01 | -0.01 | | -0.05 | 16 | 4.42 | 0.00 |
| 62 | 1.92 | 0.13 | 0.21 | 0.31 | 0.28 | -0.07 | -0.06 | 0.12 | 0.41 | 0.19 | -0.04 | 0.14 | -0.17 | -0.03 | -0.01 | | | | 17 | 4.44 | 0.00 |
| 63 | 1.91 | 0.07 | 0.20 | 0.34 | 0.28 | -0.04 | 0.04 | 0.13 | 0.33 | 0.18 | -0.04 | | 0.00 | | | -0.01 | | -0.05 | 16 | 4.49 | 0.00 |
| 64 | 1.92 | 0.09 | 0.21 | 0.33 | 0.28 | -0.06 | -0.03 | 0.12 | 0.40 | 0.19 | -0.05 | 0.01 | | | -0.01 | | 0.10 | -0.14 | 17 | 4.51 | 0.00 |
| 65 | 1.91 | 0.07 | 0.20 | 0.34 | 0.29 | -0.04 | 0.04 | 0.13 | 0.33 | 0.18 | -0.04 | | 0.00 | | 0.00 | | | -0.05 | 16 | 4.52 | 0.00 |
| 66 | 1.92 | 0.13 | 0.21 | 0.30 | 0.28 | -0.07 | -0.06 | 0.12 | 0.41 | 0.19 | -0.04 | 0.14 | -0.16 | -0.03 | | -0.01 | | | 17 | 4.54 | 0.00 |

| | Intercept | $M_w$ | $B_w$ | $F_w$ | $P_w$ | $M_w$ x $F_w$ | $M_e$ | $B_e$ | $F_e$ | $P_e$ | $M_e$ x $F_e$ | $M_w$ x $M_e$ | $M_w$ x $F_e$ | $M_w$ x $P_e$ | $B_w$ x $B_e$ | $B_w$ x $P_e$ | $F_w$ x $M_e$ | $F_w$ x $F_e$ | df | ΔAICc | Weight |
|---|---|---|---|---|---|---|---|---|---|---|---|---|---|---|---|---|---|---|---|---|---|
| 67 | 1.92 | 0.10 | 0.21 | 0.32 | 0.28 | -0.06 | -0.03 | 0.12 | 0.40 | 0.18 | -0.04 | | | | -0.01 | -0.01 | 0.10 | -0.14 | 17 | 4.58 | 0.00 |
| 68 | 1.92 | 0.09 | 0.21 | 0.33 | 0.28 | -0.06 | -0.03 | 0.12 | 0.40 | 0.19 | -0.05 | 0.01 | | | | 0.00 | 0.09 | -0.14 | 17 | 4.60 | 0.00 |
| 69 | 1.92 | 0.09 | 0.21 | 0.33 | 0.28 | -0.06 | -0.03 | 0.12 | 0.40 | 0.18 | -0.05 | | 0.01 | | -0.01 | | 0.11 | -0.15 | 17 | 4.64 | 0.00 |
| 70 | 1.92 | 0.09 | 0.21 | 0.32 | 0.28 | -0.06 | -0.03 | 0.12 | 0.40 | 0.19 | -0.04 | | 0.01 | | | 0.00 | 0.10 | -0.14 | 17 | 4.73 | 0.00 |
| 71 | 1.91 | 0.09 | 0.21 | 0.32 | 0.29 | -0.05 | 0.03 | 0.13 | 0.33 | 0.19 | -0.04 | | -0.03 | -0.03 | | | -0.02 | | 16 | 4.77 | 0.00 |
| 72 | 1.92 | 0.10 | 0.21 | 0.33 | 0.28 | -0.06 | -0.05 | 0.12 | 0.41 | 0.19 | -0.04 | 0.14 | -0.14 | -0.03 | -0.01 | | | -0.05 | 18 | 4.78 | 0.00 |
| 73 | 1.92 | 0.09 | 0.21 | 0.33 | 0.29 | -0.06 | 0.02 | 0.13 | 0.33 | 0.19 | -0.05 | | | -0.02 | 0.00 | | | | 15 | 4.80 | 0.00 |
| 74 | 1.92 | 0.10 | 0.21 | 0.33 | 0.28 | -0.06 | -0.04 | 0.12 | 0.40 | 0.19 | -0.04 | 0.20 | -0.19 | -0.02 | -0.01 | | -0.05 | | 18 | 4.80 | 0.00 |
| 75 | 1.91 | 0.09 | 0.21 | 0.33 | 0.28 | -0.06 | 0.02 | 0.13 | 0.33 | 0.19 | -0.05 | | | -0.02 | | 0.00 | | | 15 | 4.82 | 0.00 |
| 76 | 1.92 | 0.10 | 0.21 | 0.33 | 0.28 | -0.06 | -0.05 | 0.12 | 0.41 | 0.19 | -0.04 | 0.17 | -0.16 | -0.03 | | | -0.03 | -0.03 | 18 | 4.84 | 0.00 |
| 77 | 1.92 | 0.10 | 0.21 | 0.33 | 0.28 | -0.06 | -0.04 | 0.12 | 0.40 | 0.19 | -0.04 | 0.19 | -0.19 | -0.02 | | 0.00 | -0.05 | | 18 | 4.86 | 0.00 |
| 78 | 1.92 | 0.10 | 0.21 | 0.33 | 0.28 | -0.06 | -0.05 | 0.12 | 0.41 | 0.19 | -0.04 | 0.14 | -0.14 | -0.03 | | 0.00 | | -0.05 | 18 | 4.86 | 0.00 |
| 79 | 1.88 | 0.05 | 0.20 | 0.37 | 0.29 | -0.03 | | 0.13 | 0.30 | 0.17 | | | | | | | | -0.08 | 12 | 4.96 | 0.00 |
| 80 | 1.91 | 0.11 | 0.21 | 0.31 | 0.29 | -0.05 | 0.02 | 0.13 | 0.34 | 0.19 | -0.04 | | -0.04 | -0.03 | | -0.01 | | | 16 | 4.97 | 0.00 |
| 81 | 1.91 | 0.10 | 0.21 | 0.31 | 0.29 | -0.05 | 0.02 | 0.13 | 0.34 | 0.19 | -0.04 | | -0.04 | -0.03 | -0.01 | | | | 16 | 4.99 | 0.00 |
| 82 | 1.88 | 0.02 | 0.19 | 0.33 | 0.28 | | 0.07 | 0.14 | 0.32 | 0.18 | -0.04 | | | | | | -0.07 | | 13 | 5.00 | 0.00 |
| 83 | 1.91 | 0.09 | 0.21 | 0.32 | 0.29 | -0.06 | 0.04 | 0.13 | 0.32 | 0.18 | -0.05 | -0.01 | | | | -0.01 | | | 15 | 5.05 | 0.00 |
| 84 | 1.91 | 0.09 | 0.21 | 0.32 | 0.29 | -0.06 | 0.04 | 0.13 | 0.32 | 0.18 | -0.05 | -0.01 | | | -0.01 | | | | 15 | 5.06 | 0.00 |

| | Intercept | $M_w$ | $B_w$ | $F_w$ | $P_w$ | $M_w$ x $F_w$ | $M_e$ | $B_e$ | $F_e$ | $P_e$ | $M_e$ x $F_e$ | $M_w$ x $M_e$ | $M_w$ x $F_e$ | $M_w$ x $P_e$ | $B_w$ x $B_e$ | $B_w$ x $P_e$ | $F_w$ x $M_e$ | $F_w$ x $F_e$ | df | ΔAICc | Weight |
|---|---|---|---|---|---|---|---|---|---|---|---|---|---|---|---|---|---|---|---|---|---|
| 85 | 1.88 | | 0.19 | 0.35 | 0.28 | | 0.07 | 0.13 | 0.33 | 0.18 | -0.04 | | | | | -0.01 | -0.07 | | 13 | 5.10 | 0.00 |
| 86 | 1.88 | 0.08 | 0.20 | 0.34 | 0.29 | -0.03 | | 0.13 | 0.31 | 0.18 | | | -0.04 | -0.04 | | | | -0.06 | 14 | 5.12 | 0.00 |
| 87 | 1.87 | 0.02 | 0.19 | 0.35 | 0.28 | | | 0.13 | 0.33 | 0.18 | | | | -0.04 | | | | -0.11 | 12 | 5.15 | 0.00 |
| 88 | 1.88 | | 0.19 | 0.35 | 0.28 | | 0.07 | 0.14 | 0.33 | 0.18 | -0.04 | | | | 0.00 | | -0.07 | | 13 | 5.17 | 0.00 |
| 89 | 1.91 | 0.06 | 0.20 | 0.35 | 0.28 | -0.05 | 0.02 | 0.13 | 0.35 | 0.19 | -0.04 | 0.02 | | -0.02 | 0.00 | | | -0.07 | 17 | 5.17 | 0.00 |
| 90 | 1.91 | 0.06 | 0.20 | 0.35 | 0.28 | -0.05 | 0.02 | 0.13 | 0.35 | 0.19 | -0.04 | 0.02 | | -0.02 | | 0.00 | | -0.07 | 17 | 5.18 | 0.00 |
| 91 | 1.92 | 0.09 | 0.21 | 0.33 | 0.29 | -0.06 | 0.03 | 0.13 | 0.32 | 0.18 | -0.05 | | | | -0.01 | -0.01 | | | 15 | 5.27 | 0.00 |
| 92 | 1.92 | 0.12 | 0.21 | 0.30 | 0.28 | -0.07 | -0.05 | 0.12 | 0.40 | 0.19 | -0.05 | 0.14 | -0.16 | | -0.02 | -0.01 | | | 17 | 5.39 | 0.00 |
| 93 | 1.88 | 0.03 | 0.19 | 0.32 | 0.28 | | 0.03 | 0.13 | 0.37 | 0.18 | -0.04 | | | | | -0.01 | | -0.08 | 14 | 5.40 | 0.00 |
| 94 | 1.91 | 0.07 | 0.20 | 0.34 | 0.28 | -0.04 | 0.03 | 0.13 | 0.34 | 0.19 | -0.04 | | | -0.03 | -0.01 | -0.01 | | -0.05 | 17 | 5.45 | 0.00 |
| 95 | 1.91 | 0.07 | 0.21 | 0.34 | 0.28 | -0.05 | 0.05 | 0.13 | 0.31 | 0.19 | -0.04 | | | -0.02 | | 0.00 | -0.03 | | 16 | 5.45 | 0.00 |
| 96 | 1.89 | 0.03 | 0.19 | 0.32 | 0.28 | | 0.02 | 0.13 | 0.37 | 0.18 | -0.04 | | | | -0.01 | | | -0.08 | 14 | 5.46 | 0.00 |
| 97 | 1.91 | 0.08 | 0.20 | 0.34 | 0.28 | -0.04 | 0.03 | 0.13 | 0.34 | 0.19 | -0.04 | | -0.01 | -0.03 | | 0.00 | | -0.05 | 17 | 5.46 | 0.00 |
| 98 | 1.91 | 0.07 | 0.21 | 0.34 | 0.29 | -0.05 | 0.05 | 0.13 | 0.31 | 0.19 | -0.04 | | | -0.02 | 0.00 | | -0.03 | | 16 | 5.47 | 0.00 |
| 99 | 1.91 | 0.07 | 0.21 | 0.34 | 0.29 | -0.05 | 0.05 | 0.13 | 0.31 | 0.19 | -0.04 | 0.00 | | -0.02 | | | -0.04 | | 16 | 5.47 | 0.00 |
| 100 | 1.91 | 0.07 | 0.20 | 0.34 | 0.28 | -0.04 | 0.03 | 0.13 | 0.34 | 0.19 | -0.04 | | -0.01 | -0.03 | 0.00 | | | -0.05 | 17 | 5.47 | 0.00 |
| 101 | 1.89 | 0.03 | 0.19 | 0.32 | 0.28 | | 0.02 | 0.13 | 0.38 | 0.18 | -0.04 | | | | | | 0.01 | -0.09 | 14 | 5.48 | 0.00 |
| 102 | 1.89 | 0.03 | 0.19 | 0.32 | 0.28 | | 0.02 | 0.13 | 0.37 | 0.18 | -0.04 | | -0.01 | | | | | -0.07 | 14 | 5.48 | 0.00 |

| | Intercept | $M_w$ | $B_w$ | $F_w$ | $P_w$ | $M_w$ x $F_w$ | $M_e$ | $B_e$ | $F_e$ | $P_e$ | $M_e$ x $F_e$ | $M_w$ x $M_e$ | $M_w$ x $F_e$ | $M_w$ x $P_e$ | $B_w$ x $B_e$ | $B_w$ x $P_e$ | $F_w$ x $M_e$ | $F_w$ x $F_e$ | df | ΔAICc | Weight |
|---|---|---|---|---|---|---|---|---|---|---|---|---|---|---|---|---|---|---|---|---|---|
| 103 | 1.88 | 0.12 | 0.21 | 0.31 | 0.29 | -0.04 | | 0.12 | 0.30 | 0.18 | | | -0.07 | -0.04 | | | | | 13 | 5.49 | 0.00 |
| 104 | 1.89 | 0.03 | 0.19 | 0.32 | 0.28 | | 0.02 | 0.13 | 0.37 | 0.18 | -0.04 | 0.00 | | | | | | -0.08 | 14 | 5.50 | 0.00 |
| 105 | 1.92 | 0.10 | 0.21 | 0.32 | 0.28 | -0.06 | -0.05 | 0.12 | 0.41 | 0.19 | -0.04 | | | -0.03 | -0.01 | -0.01 | 0.10 | -0.15 | 18 | 5.64 | 0.00 |
| 106 | 1.89 | | 0.19 | 0.34 | 0.28 | | 0.02 | 0.13 | 0.38 | 0.18 | -0.04 | | | | -0.01 | -0.01 | | -0.08 | 14 | 5.64 | 0.00 |
| 107 | 1.92 | 0.10 | 0.21 | 0.32 | 0.28 | -0.06 | -0.05 | 0.12 | 0.41 | 0.19 | -0.04 | | 0.00 | -0.03 | -0.01 | | 0.10 | -0.15 | 18 | 5.67 | 0.00 |
| 108 | 1.92 | 0.09 | 0.21 | 0.33 | 0.28 | -0.06 | -0.05 | 0.12 | 0.41 | 0.19 | -0.04 | 0.00 | | -0.03 | -0.01 | | 0.10 | -0.15 | 18 | 5.67 | 0.00 |
| 109 | 1.92 | 0.09 | 0.21 | 0.33 | 0.28 | -0.06 | -0.03 | 0.12 | 0.40 | 0.19 | -0.04 | 0.20 | -0.19 | | -0.01 | -0.01 | -0.05 | | 18 | 5.69 | 0.00 |
| 110 | 1.87 | 0.05 | 0.19 | 0.32 | 0.28 | | | 0.13 | 0.34 | 0.18 | | | -0.04 | -0.04 | | | | -0.07 | 13 | 5.75 | 0.00 |
| 111 | 1.92 | 0.10 | 0.21 | 0.32 | 0.28 | -0.06 | -0.04 | 0.12 | 0.41 | 0.19 | -0.04 | | 0.00 | -0.03 | | 0.00 | 0.10 | -0.14 | 18 | 5.76 | 0.00 |
| 112 | 1.92 | 0.09 | 0.21 | 0.33 | 0.28 | -0.06 | -0.04 | 0.12 | 0.41 | 0.19 | -0.04 | 0.15 | -0.14 | | -0.01 | -0.01 | | -0.05 | 18 | 5.76 | 0.00 |
| 113 | 1.92 | 0.09 | 0.21 | 0.33 | 0.28 | -0.06 | -0.04 | 0.12 | 0.41 | 0.19 | -0.04 | 0.00 | | -0.03 | | 0.00 | 0.10 | -0.15 | 18 | 5.76 | 0.00 |
| 114 | 1.89 | | 0.19 | 0.34 | 0.28 | | 0.01 | 0.13 | 0.38 | 0.18 | -0.04 | | | | -0.01 | | 0.01 | -0.08 | 14 | 5.77 | 0.00 |
| 115 | 1.88 | 0.04 | 0.19 | 0.31 | 0.28 | | 0.00 | 0.13 | 0.39 | 0.19 | -0.03 | | -0.02 | -0.03 | | | | -0.07 | 15 | 5.78 | 0.00 |
| 116 | 1.88 | | 0.19 | 0.34 | 0.28 | | 0.02 | 0.13 | 0.38 | 0.18 | -0.04 | | | | | -0.01 | 0.00 | -0.08 | 14 | 5.79 | 0.00 |
| 117 | 1.92 | 0.09 | 0.21 | 0.34 | 0.28 | -0.06 | -0.04 | 0.12 | 0.40 | 0.19 | -0.04 | 0.19 | -0.17 | | -0.01 | | -0.04 | -0.02 | 18 | 5.81 | 0.00 |
| 118 | 1.88 | 0.02 | 0.19 | 0.33 | 0.28 | | 0.06 | 0.14 | 0.33 | 0.19 | -0.03 | | | -0.03 | | | -0.07 | | 14 | 5.83 | 0.00 |
| 119 | 1.92 | 0.09 | 0.21 | 0.33 | 0.28 | -0.06 | -0.03 | 0.12 | 0.40 | 0.19 | -0.04 | 0.19 | -0.17 | | | -0.01 | -0.04 | -0.01 | 18 | 5.88 | 0.00 |
| 120 | 1.89 | 0.04 | 0.19 | 0.32 | 0.28 | | 0.01 | 0.13 | 0.38 | 0.19 | -0.03 | -0.01 | | -0.03 | | | | -0.08 | 15 | 5.95 | 0.00 |

| | Intercept | $M_w$ | $B_w$ | $F_w$ | $P_w$ | $M_w$ x $F_w$ | $M_e$ | $B_e$ | $F_e$ | $P_e$ | $M_e$ x $F_e$ | $M_w$ x $M_e$ | $M_w$ x $F_e$ | $M_w$ x $P_e$ | $B_w$ x $B_e$ | $B_w$ x $P_e$ | $F_w$ x $M_e$ | $F_w$ x $F_e$ | df | ΔAICc | Weight |
|---|---|---|---|---|---|---|---|---|---|---|---|---|---|---|---|---|---|---|---|---|---|
| 121 | 1.89 | 0.03 | 0.19 | 0.32 | 0.28 | | 0.01 | 0.13 | 0.38 | 0.19 | -0.03 | | | -0.03 | | -0.01 | | -0.09 | 15 | 6.00 | 0.00 |
| 122 | 1.89 | 0.03 | 0.19 | 0.33 | 0.28 | | 0.01 | 0.13 | 0.38 | 0.19 | -0.03 | | | -0.03 | 0.00 | | | -0.09 | 15 | 6.01 | 0.00 |
| 123 | 1.89 | 0.03 | 0.19 | 0.32 | 0.28 | | 0.00 | 0.13 | 0.39 | 0.19 | -0.03 | | | -0.03 | | | 0.01 | -0.10 | 15 | 6.01 | 0.00 |
| 124 | 1.91 | 0.06 | 0.20 | 0.35 | 0.28 | -0.05 | 0.03 | 0.13 | 0.34 | 0.18 | -0.05 | 0.03 | | | -0.01 | -0.01 | | -0.07 | 17 | 6.05 | 0.00 |
| 125 | 1.91 | 0.09 | 0.21 | 0.32 | 0.29 | -0.05 | 0.05 | 0.13 | 0.32 | 0.18 | -0.04 | | -0.02 | | | -0.01 | -0.02 | | 16 | 6.08 | 0.00 |
| 126 | 1.87 | 0.02 | 0.19 | 0.35 | 0.29 | | | 0.13 | 0.33 | 0.17 | | | | | | | | -0.10 | 11 | 6.11 | 0.00 |
| 127 | 1.91 | 0.08 | 0.21 | 0.33 | 0.29 | -0.05 | 0.05 | 0.13 | 0.32 | 0.18 | -0.04 | | -0.02 | | 0.00 | | -0.02 | | 16 | 6.14 | 0.00 |
| 128 | 1.86 | | 0.19 | 0.37 | 0.29 | | | 0.13 | 0.33 | 0.17 | | | | | | -0.01 | | -0.10 | 11 | 6.17 | 0.00 |
| 129 | 1.87 | | 0.19 | 0.37 | 0.29 | | -0.02 | 0.13 | 0.35 | 0.17 | | | | | | | | -0.10 | 11 | 6.18 | 0.00 |
| 130 | 1.87 | | 0.19 | 0.37 | 0.29 | | | 0.13 | 0.33 | 0.17 | | | | | -0.01 | | | -0.10 | 11 | 6.19 | 0.00 |
| 131 | 1.91 | 0.10 | 0.21 | 0.31 | 0.29 | -0.05 | 0.04 | 0.13 | 0.33 | 0.18 | -0.04 | | -0.03 | | -0.01 | -0.01 | | | 16 | 6.20 | 0.00 |
| 132 | 1.91 | 0.06 | 0.21 | 0.34 | 0.29 | -0.05 | 0.06 | 0.13 | 0.31 | 0.18 | -0.05 | 0.01 | | | | -0.01 | -0.04 | | 16 | 6.30 | 0.00 |
| 133 | 1.91 | 0.06 | 0.21 | 0.35 | 0.29 | -0.05 | 0.06 | 0.13 | 0.31 | 0.18 | -0.05 | 0.01 | | | 0.00 | | -0.04 | | 16 | 6.32 | 0.00 |
| 134 | 1.91 | 0.07 | 0.20 | 0.33 | 0.29 | -0.05 | 0.06 | 0.13 | 0.31 | 0.18 | -0.05 | | | | -0.01 | -0.01 | -0.03 | | 16 | 6.34 | 0.00 |
| 135 | 1.88 | 0.05 | 0.20 | 0.37 | 0.29 | -0.03 | -0.02 | 0.13 | 0.32 | 0.18 | | | | -0.04 | | | | -0.08 | 14 | 6.36 | 0.00 |
| 136 | 1.91 | 0.10 | 0.21 | 0.32 | 0.29 | -0.05 | 0.03 | 0.13 | 0.32 | 0.19 | -0.04 | -0.02 | | -0.02 | | -0.01 | | | 16 | 6.40 | 0.00 |
| 137 | 1.91 | 0.09 | 0.21 | 0.32 | 0.29 | -0.05 | 0.03 | 0.13 | 0.32 | 0.19 | -0.04 | -0.02 | | -0.02 | 0.00 | | | | 16 | 6.41 | 0.00 |
| 138 | 1.88 | 0.05 | 0.20 | 0.37 | 0.29 | -0.03 | | 0.13 | 0.30 | 0.18 | | | | -0.04 | | 0.00 | | -0.08 | 14 | 6.42 | 0.00 |

| | Intercept | $M_w$ | $B_w$ | $F_w$ | $P_w$ | $M_w$ x $F_w$ | $M_e$ | $B_e$ | $F_e$ | $P_e$ | $M_e$ x $F_e$ | $M_w$ x $M_e$ | $M_w$ x $F_e$ | $M_w$ x $P_e$ | $B_w$ x $B_e$ | $B_w$ x $P_e$ | $F_w$ x $M_e$ | $F_w$ x $F_e$ | df | ΔAICc | Weight |
|---|---|---|---|---|---|---|---|---|---|---|---|---|---|---|---|---|---|---|---|---|---|
| 139 | 1.88 | 0.05 | 0.20 | 0.37 | 0.29 | -0.03 | | 0.13 | 0.30 | 0.18 | | | | -0.04 | 0.00 | | | -0.08 | 14 | 6.43 | 0.00 |
| 140 | 1.88 | 0.07 | 0.20 | 0.35 | 0.29 | -0.03 | | 0.13 | 0.31 | 0.17 | | | -0.03 | | | | | -0.06 | 13 | 6.50 | 0.00 |
| 141 | 1.92 | 0.13 | 0.21 | 0.30 | 0.28 | -0.07 | -0.06 | 0.12 | 0.41 | 0.19 | -0.04 | 0.14 | -0.17 | -0.03 | -0.02 | -0.01 | | | 18 | 6.62 | 0.00 |
| 142 | 1.91 | 0.07 | 0.20 | 0.34 | 0.28 | -0.04 | 0.04 | 0.13 | 0.33 | 0.18 | -0.04 | | 0.00 | | -0.01 | -0.01 | | -0.05 | 17 | 6.72 | 0.00 |
| 143 | 1.92 | 0.09 | 0.21 | 0.33 | 0.28 | -0.06 | -0.03 | 0.12 | 0.40 | 0.18 | -0.05 | 0.01 | | | -0.01 | -0.01 | 0.10 | -0.14 | 18 | 6.78 | 0.00 |
| 144 | 1.88 | 0.04 | 0.19 | 0.32 | 0.29 | | 0.06 | 0.13 | 0.34 | 0.18 | -0.04 | | -0.02 | | | | -0.05 | | 14 | 6.86 | 0.00 |
| 145 | 1.92 | 0.09 | 0.21 | 0.32 | 0.28 | -0.06 | -0.03 | 0.12 | 0.40 | 0.18 | -0.04 | | 0.00 | | -0.01 | -0.01 | 0.11 | -0.15 | 18 | 6.89 | 0.00 |
| 146 | 1.89 | 0.10 | 0.21 | 0.35 | 0.28 | -0.05 | -0.08 | 0.12 | 0.39 | 0.19 | | 0.19 | -0.21 | -0.04 | | | -0.06 | | 16 | 6.93 | 0.00 |
| 147 | 1.88 | 0.10 | 0.21 | 0.32 | 0.29 | -0.04 | | 0.12 | 0.29 | 0.17 | | | -0.06 | | | | | | 12 | 6.95 | 0.00 |

**Table S5:** Parameter estimates, 95% confidence intervals (CI), and z-values for final averaged model describing the effects of key variance components on mean lifetime reproductive success.

| Component | Estimate | Lower CI | Upper CI | z value |
|---|---|---|---|---|
| $M_w$ | 0.08 | -0.03 | 0.19 | 1.29 |
| $B_w$ | 0.20 | 0.16 | 0.25 | 8.39 |
| $F_w$ | 0.33 | 0.23 | 0.44 | 6.18 |
| $P_w$ | 0.28 | 0.23 | 0.34 | 9.62 |
| $M_w$ x $F_w$ | -0.05 | -0.10 | -0.01 | 1.68 |
| $M_e$ | 0.00 | -0.19 | 0.19 | 0.04 |
| $B_e$ | 0.13 | 0.06 | 0.19 | 3.97 |
| $F_e$ | 0.36 | 0.18 | 0.54 | 3.96 |
| $P_e$ | 0.19 | 0.12 | 0.25 | 5.79 |
| $M_e$ x $F_e$ | -0.04 | -0.07 | -0.01 | 2.36 |
| $M_w$ x $M_e$ | 0.04 | -0.11 | 0.32 | 0.44 |
| $M_w$ x $F_e$ | -0.04 | -0.30 | 0.11 | 0.46 |
| $M_w$ x $P_e$ | -0.01 | -0.07 | 0.02 | 0.48 |
| $B_w$ x $B_e$ | 0.00 | -0.06 | 0.05 | 0.12 |
| $B_w$ x $P_e$ | 0.00 | -0.06 | 0.05 | 0.09 |
| $F_w$ x $M_e$ | 0.01 | -0.16 | 0.22 | 0.19 |
| $F_w$ x $F_e$ | -0.05 | -0.22 | 0.05 | 0.79 |
